# Assessing single-component gene drive systems in the mosquito *Aedes aegypti* via single generation crosses and modeling

**DOI:** 10.1101/2021.12.08.471839

**Authors:** William Reid, Adeline E Williams, Irma Sanchez-Vargas, Jingyi Lin, Rucsanda Juncu, Ken E Olson, Alexander WE Franz

**Affiliations:** Department of Veterinary Pathobiology, University of Missouri, Columbia, MO 65211, USA; Department of Microbiology, Immunology, and Pathology, Colorado State University, Fort Collins, CO 80523, USA; Laboratory of Malaria and Vector Research, National Institute of Allergy and Infectious Diseases, National Institutes of Health, Rockville, MD, 20852, USA

**Keywords:** gene drive blocking indels, arbovirus, CRISPR, genomic position effect, population replacement

## Abstract

The yellow fever mosquito *Aedes aegypti* is a major vector of arthropod-borne viruses, including dengue, chikungunya, and Zika. A novel approach to mitigate arboviral infections is to generate mosquitoes refractory to infection by overexpressing antiviral effector molecules. Such an approach requires a mechanism to spread these antiviral effectors through a population, for example, by using CRISPR/Cas9-based gene drive (GD) systems. Critical to the design of a single-locus autonomous GD is that the selected genomic locus be amenable to both GD and appropriate expression of the antiviral effector. In our study, we used reverse engineering to target two intergenic genomic loci, which had previously shown to be highly permissive for antiviral effector gene expression, and we further investigated the use of three promoters (*nanos, β2-tubulin,* or *zpg*) for Cas9 expression. We then quantified the accrual of insertions or deletions (indels) after single generation crossings, measured maternal effects, and assessed fitness costs associated with the various transgenic lines to model the rate of GD fixation. Overall, MGDrivE modeling suggested that when an autonomous GD is placed into an intergenic locus, the GD system will eventually be blocked by the accrual of GD blocking resistance alleles and ultimately be lost in the population. Moreover, while genomic locus and promoter selection were critically important for the initial establishment of the autonomous GD, it was the fitness of the GD line that most strongly influenced the persistence of the GD in the simulated population. As such, we propose that when autonomous CRISPR/Cas9 based GD systems are anchored in an intergenic locus, they temporarily result in a strong population replacement effect, but as GD-blocking indels accrue, the GD becomes exhausted due to the fixation of CRISPR resistance alleles.

**Significance statement:** For the purpose of population replacement, CRISPR/Cas9 based gene drives (GD) have been developed in *Anopheles* spp. and split GDs have been developed in *Ae. aegypti.* In our study, we developed autonomous GD in *Ae. aegypti* and positioned the drives in intergenic loci ideal for the expression of antiviral effector genes. Our results suggest that when the GD is placed into an intergenic locus, there is rapid introgression of the GD resulting in a transient population replacement followed by loss of the drive as resistance alleles accrue. Fitness of the transgenic lines and maternal deposition of CRISPR/Cas9 components were the major contributing factors affecting the perseverance of the GD in our population models.

## INTRODUCTION

The yellow fever mosquito *Aedes aegypti* is the principal vector of arthropod-borne viruses (arboviruses) such as dengue, yellow fever, chikungunya, and Zika in tropical regions of the world (1–3). Presently, the major strategy to control *Ae. aegypti* populations relies on the use of *Bacillus thuringiensis* var *israelensis* (*Bti*) and chemical insecticides; however, these approaches have led to multiple mechanisms of insecticide resistance, warranting the ongoing quest for alternative methods of control (4–5). Novel approaches, including the incompatible insect technique (IIT) using the intracellular parasite *Wolbachia* (6, 7) and transgenic mosquitoes containing dominant lethal transgenes (RIDL, fsRIDL), have been tested in the field (8, 9). Other promising technologies, including precision-guided sterile insect technique (pgSIT) have been developed for the purposes of localized, confinable management of *Ae. aegypti* populations (10, 11). Another genetic control strategy, termed population replacement, aims at spreading an antipathogen (*i.e.*, antiviral) effector through a targeted mosquito population. Antiviral effectors, specifically those blocking dengue and Zika viruses in *Ae. aegypti*, have been previously developed and tested (12–17). Linking an autonomous gene drive (GD) system to the antiviral effector could lead to super-Mendelian inheritance of the transgene within the targeted mosquito population (18). As a consequence, the proportion of the population harboring the effectors would be increased, resulting in the emergence of mosquitoes refractory to arbovirus transmission. For such a broad-scale population replacement approach, the GD system needs to overcome several limitations, including: 1) the antiviral effector must provide sufficient efficacy to render the targeted population refractory, 2) the inheritance of the GD must be greater than any fitness cost associated with the GD, and 3) the GD must be able to outpace the development of GD resistant alleles. GD systems utilizing a homing endonuclease gene (HEG) based approach were initially proposed in 2003 and subsequently demonstrated in 2008 in *Anopheles gambiae* as a fully synthetic homing GD using the DNA cleaving intron-encoded endonuclease I-SceI (19). The CRISPR/Cas9 system has been modified to allow for HEG-based GDs in *Drosophila melanogaster* (20), along with several mosquito species including *Anopheles stephensi* (21, 22), *An. gambiae* (23), and *Ae. aegypti* (10, 24). The HEG-based GD developed in *Ae. aegypti* inserted itself into the *white* gene on chromosome 1 and contained a single guide RNA (sgRNA)-expressing cassette that targeted the locus, along with an eGFP eye marker as cargo (10).

Highly invasive GD systems are thought to carry substantial environmental risk since they are not designed to self-eliminate or be confinable to a region. However, several systems have been developed that would allow for their recall, including systems that destroy or overwrite the GD, and more recently, a “biodegradable” self-eliminating GD system (25–28). Split-GD systems have been developed for *Ae. aegypti*; by design – these GD systems are modelled to be self-limiting or self-extinguishing (10, 24). Five different *Ae. aegypti* lines, each of which expressed Cas9 under the control of a different promoter (*i.e*., *exuperantia*, *4nitro*, *trunk*, *nup50*, and *PUb*) were used to drive *in trans* an sgRNA-expressing cassette positioned in the *white* gene (10, 29). The work by Li *et al* (2020) demonstrated an efficient split GD system that was designed to be confinable (10). Of major concern to the development of an autonomous GD system with an associated genetically linked antipathogen effector is that the expression patterns for the HEG-based GD and the antipathogen effector are spatio-temporally distinct from each other. That is, the HEG-based GD is ideally active in the germline during early gametogenesis while the antipathogen effector is ideally expressed in tissues associated with the pathogen in the mosquitoes, such as in the female midgut. Therefore, the genomic locus for an autonomous CRISPR/Cas9 GD must allow for both the expression of the GD as well as the antipathogen effector. Multiple viral effectors have been developed and tested in *Ae. aegypti* (30), with these effectors all being incorporated into the genome in a quasi-random fashion using transposons. While multiple genomic loci have been identified to be suitable for transgene expression in *Ae. aegypti*, two of our previous studies have identified genomic loci that reliably allow for high levels of gene expression in the female midgut following ingestion of a blood meal (13, 31). In our studies, midgut-specific transgenes were placed under control of the *carboxypeptidase A* promoter and identified two genomic loci that exhibited strong transgene expression: ‘Carb 109’ (C109) and ‘TIMP-P4’ (T4) (13, 31). In addition, the Carb109 locus was found to be highly stable for more than 50 generations for a dengue virus type 2 (DENV2)-targeting inverted-repeat effector and represents an ideal genomic locus for the insertion and expression of antiviral effectors. In our study, we took a reverse engineering approach to design, build, and test autonomous CRISPR/Cas9-based GD systems that position the GD at intergenic loci known to allow for efficient transgene expression in the midgut and investigated three promoters/3’-UTR for the expression of the Cas9 nuclease. Following establishment, we characterized the GD along with fitness parameters of the GD harboring lines and modeled how they would behave as a single release in a confined area. Overall, our study demonstrates the effects of placing an autonomous GD system in an intergenic genomic locus and highlights the many variables that ultimately affect transgenic population fixation in the context of GD performance.

## MATERIALS AND METHODS

### Generation of cDNA constructs

The constructs used for establishing the GD lines were prepared in three sequential steps: 1) an ‘empty’ gene drive destination vector containing the homology arms corresponding to the destination locus (*i.e.*: TIMP-P4 or Carb109); 2) the NLS-Cas9 gene from the pHsp70 Cas9 plasmid (Addgene plasmid #46294 (32)) connected to the promoters and 3’-UTRs from the *β2tubulin* (AAEL019894), *nanos* (AAEL012107), and *innexin-4* (*zpg*, AAEL006726) genes, respectively, 3) the respective sgRNA targeting either the TIMP-P4 or the Carb109 locus under control of the U6:3 snRNA promoter (AAEL017774).

#### Construction of ‘empty’ destination vectors for the GD cassettes

The ‘empty’ destination vector for the TIMP-P4 locus (Chr2:32138225) was constructed as previously described for plasmid AeaeCFPT4 (**Table S6**) with the exception that the eCFP marker was replaced with that of mCherry as a two-fragment Gibson assembly (17). The ORF of mCherry was amplified using primers BR-100 and BR-101, while primers BR-98 and BR-99 were used to amplify the full backbone of the destination vector minus the eCFP marker (**Table S5**). The two PCR fragments were then assembled using the HiFi Gibson Assembly kit from NEB (Waltham, MA) following the manufacturer’s instructions. The ‘empty’ destination vector for the Carb109 locus (3: 409699138) was constructed by first amplifying the 459 bp upstream homology arm (genomic positions AaegL5_3:409698681-409699140) using primers BR-368 and BR-122, followed by insertion of the amplicon into pBluescript using *Kpn*I and *Xho*I. The 541 bp downstream homology arm (genomic positions AaegL5_3: 3:409699139-409699680) was then amplified using primers BR-113 and BR-364, followed by insertion of the amplicon into the previous intermediate plasmid using *Xho*I and *Sac*II. Finally, the 3xP3-mCherry-SV40 cassette was amplified from the empty TIMP-P4 destination vector using primers BR-54 and BR-55 and cloned into the destination vector using *Xho*I followed by screening for orientation using primers BR-60 and BR-54.

#### Construction of the β2tubulinCas9 cassette

The basic components of the β2tubulinCas9 construct were assembled in pUC19 in three steps. First, the *β2tubulin* promoter/5’-UTR (AaegL5_2: 326339137-326338047) fragment was amplified from the *Ae. aegypti* genome (HWE strain) using primers BR-32 and BR-33 and cloned into pUC19 using *Hind*III and *Pst*I, which introduced an *Xho*I site upstream of the promoter and an *Nco*I site at the end of the 5’-UTR of *β2tubulin*. The 4272 bp ORF of Cas9 was then amplified from pHsp70 Cas9 plasmid (Addgene plasmid #46294 (32)) using primers BR-34 and BR-35 and inserted into the plasmid vector using *Nco*I and *Sal*I. Finally, the 3’-UTR fragment of *β2tubulin* (AaegL5_2:326336649-326336454) was amplified using BR-28 and BR-29 and inserted into the assembly plasmid vector using *Sal*I and *Xba*I. The entire cassette was then removable from pUC19 using *Xho*I and *Xba*I for cloning into the TIMP-P4 or Carb109 destination vectors as either *Xho*I/*Xba*I or *Xho*I/blunted insertions.

#### Construction of the nanosCas9 cassette

The 1159 bp *nanos* promoter fragment (AaegL5_1:228706-229865) was amplified from the HWE strain of *Ae. aegypti* using primers BR-40 and BR-41 and cloned into pBluescript using *Xho*I followed by screening for orientation using primers BR-99 and BR-41. The 4272 bp ORF of Cas9 was then amplified from the pHsp70 Cas9 plasmid (Addgene plasmid #46294 (32)) using primers BR-34 and BR-35 and inserted into pBluescript downstream of the *nanos* promoter fragment using a *Nco*I and *Sal*I digest for the PCR product and a *Pci*I/*Sal*I digest for the plasmid vector. Finally, the 594 bp *nanos* 3’-UTR fragment (AaegL5_1:240330-240924) was amplified from the *Ae. aegypti* genome (HWE strain) using primers BR-42 and BR-43 and cloned into the assembly plasmid vector downstream of the Cas9 ORF using *Stu*I and *Nhe*I.

#### Construction of Carb109-zpgCas9^GD^

The 1729 bp *zpg* promoter fragment (AaegL5_2:84862322-84863150) was amplified from genomic DNA of the HWE strain of *Ae. aegypti* using primers BR-655 and BR-656. The promoter region for the *zpg* gene in the HWE strain contained a 144 bp deletion (positions AaegL5_2:84863229-848632372), which was 825 bp upstream of the +1 ATG of the *zpg* ORF. The PCR product containing the promoter region was digested with *Xho*I and *Xba*I and cloned into pBluescript. In addition, primer BR-655 was internally tagged with a *Bbs*I site to allow for downstream scarless introduction of Cas9. The 1357 bp 3’-UTR of the *zpg* gene (AaegL5_2:84865497-84866853) was amplified from *Ae. aegypti* (HWE strain) genomic DNA using primers BR-660 and BR-683 and cloned into the vector containing the *zpg* promoter using *Pst*I and *Spe*I. In addition, primer BR-660 was internally tagged with an inverted *Bbs*I site to allow for downstream scarless introduction of Cas9. Subsequently, the 4272 bp ORF of Cas9 was amplified from pHsp70 Cas9 plasmid (Addgene plasmid #46294 (32)) using primers BR-666 and BR-667 and cloned into the vector between the *zpg* promoter and the 3’-UTR for *zpg* using *Bbs*I for scarless, directional cloning.

#### Assembly of the Carb109-3xP3-eCFP eye marker vector

The Carb109-3xP3-eCFP vector was constructed by replacing the mCherry ORF from the Carb109 destination vector with eCFP. The eCFP ORF was amplified from plasmid AeaeCFPT4 using primers BR-100 and BR-101 (17). The mCherry-deleted backbone from the empty Carb109 destination vector was amplified using BR-98 and BR-99. The two PCR fragments were then assembled using the HiFi Gibson Assembly kit from NEB (Waltham, MA) following the manufacturer’s instructions.

#### Assembly of the Pol-III promoter/chiRNA ‘empty’ vector

The active U6 promoter from Konet *et al*., 2007 (33) was synthesized as a gBlock (IDT, Corralville, IA) starting with the promoter region for AAEL017774 (AaegL5_3:382122755-382123154, minus strand) with a G to A substitution at the −6 position relative to the TATA box in order to domesticate a *Bbs*I restriction site and allow for Golden-Gate cloning (34). The sgRNA scaffold from Dang *et al*. (2015) (35) was placed downstream of the promoter and separated by two inverted *Bbs*I sites oriented such that the generated 5’ overhang regions were present within the promoter and chiRNA respectively (36). Primers tagged with *Sac*II restriction sites were used to PCR amplify the full gBlock cDNA, which was then cloned into pBluescript using *Sac*II. Subsequently, the sgRNA programming for all gRNA was prepared by digesting the empty Pol III promoter/chiRNA vector with *Bbs*I followed by ligation of the respective adapter primers (BR-360 and BR-361 for the TIMP-P4 locus, BR-362 and BR-363 for the Carb109 locus). Adapters were unphosphorylated and tagged with 5’-AAAT for the protospacer sequence (PAM-sense strand) and 5’-AAAC for the corresponding reverse complement (PAM antisense strand), following the approach outlined in Gokcezade *et al*., 2014 (36). The respective programmed U6-sgRNA cassettes were then amplified using BR-350 and BR-351 and cloned into the destination vectors using *Sac*II followed by screening for orientation using the appropriate forward adapter (BR-360 for TIMP-P4 and BR-362 for Carb109) and M13R primers.

#### Assembly of the pAeT7ku70 plasmid for dsRNA production

A total of 10 fourth instar *Ae. aegypti* (HWE strain) larvae were collected, flash frozen on dry ice and then extracted for total RNA using TRIzol reagent (Carlsbad, CA). First strand cDNA synthesis was performed using the Protoscript cDNA kit (NEB, Waltham, MA) following the manufacturer’s instructions. The ku70 dsRNA template was then amplified from the cDNA using primers BR-44 and BR-45, which tagged both ends of the PCR product with the T7 promoter (37).

Subsequently, the T7-tagged PCR product was amplified using *Hind*III-tagged T7 (BR-80) and cloned into pUC19 using *Hind*III.

#### Final assemblies of GD constructs

The final assemblies of the gene drive constructs were prepared using a three-step cloning strategy. Cas9 cassettes for the TIMP-P4 or the Carb109 locus were cloned into their respective destination vectors using *Xho*I/*Sac*II, followed by the addition of the mCherry fluorescent marker under control of the 3xP3 synthetic promoter (38) using *Xho*I, and finally the addition of the appropriate U6 promoter plus sgRNA using *Sac*II. The schematic for our GD constructs is presented in **Figure S2**, and the primers used in our study are listed in **Table S5**. The final sequences for the constructs are available at NCBI as outlined in **Table S6**.

### Generation of transgenic lines

#### Mosquito rearing

The HWE strain of *Ae. aegypti* was used for the generation of all transgenic lines in our study. The larval stages were reared in deionized water at 28°C with a 12:12 (L:D) photoperiod and fed a diet of Tetramin Tropical flakes (Tetra Spectrum Brands Pet, LLC, Blacksburg, VA). Pupae were transferred to plastic containers and allowed to emerge in 18 cubic-inch cages and provided with deionized water and raisins as food sources.

#### CRISPR/Cas9 target site identification

For our initial testing, we first assessed the activities of six sgRNAs for activity proximal to the Carb109 genome locus, while the active sgRNA for the TIMP-P4 genome locus was already reported (17). To accomplish this, we prepared all injection mix material so that the final concentrations of Cas9-NLS protein was 300 ng/µL and the concentration of sgRNA was 80 ng/µL. Then, three sets of approximately 100 pre-blastoderm embryos were injected with injection mix and incubated overnight (18 h – 24 h) at 27°C before being homogenized and extracted for gDNA into each of three pools. The genomic sequence of the Carb109 locus was then PCR amplified using primers BR-60 and BR-65, and the resulting amplicons gel-purified using the Zymo Clean and Concentrator kit (Zymo Research, San Diego, CA). PCR products were subsequently sequenced at the University of Missouri Genomics Technology Core and analyzed visually for trace decay at the predicted CRISPR/Cas9 cleavage site. All products were sequenced in both directions to confirm the correct site of sequence decay.

#### Harvest and microinjection of embryos

Cages of 7-10 day-old mosquitoes were provided with a blood meal (defibrinated sheep blood, Colorado Serum, Denver, CO) and maintained at 28°C under a 12:12 (L:D) photoperiod for four days prior to injection. Groups of 15-20 hyper-gravid females were then mouth aspirated into an aluminum foil-wrapped 50 mL conical tube containing two 2.5 x 0.5 cm strips of moistened, overlapping Whatman #1 filter paper and allowed to oviposit for 20 min. Early embryos were then aligned over a period of 20 to 30 min using a fine spotter paint brush such that all posterior ends of the embryos faced the same direction. Aligned embryos were then transferred to double-faced tape (Scotch brand, 3M Columbia, MO) affixed to a plastic coverslip (Fisher Scientific, Waltham, MA) and covered with a layer of Halocarbon 27 oil (Sigma-Aldrich, St. Louis, MO) prior to the visible onset of melanization. Embryos were then injected with injection mix using a pulled and beveled micropipette capillary tube (Sutter Instruments, Novato, CA) connected to a Femtojet 5247 air compressor (Eppendorf, Germany) at 650 psi positive pressure and 100 psi backpressure. Immediately following injection, the Halocarbon 27 oil was gently washed from the embryos with deionized water, and the embryos were allowed to age in a humid chamber for a minimum of 2 h prior to transfer to a 500 mL plastic beaker lined with 5-10 layers of moistened Kimwipes. Injected embryos (herein referred to as G_0_) were then maintained in the humid plastic beaker for 6 days, transferred to a small plastic cup filled with deionized water and allowed to hatch out. G_0_ survivors were reared as described above and backcrossed to non-transgenic HWE. For the backcrossing, male G_0_ individuals were individually provided with 5-10 virgin females and allowed to mate for 3-5 days then pooled into larger cartons using 20 of the smaller cartons for each large carton. Female G_0_ were collected into large cartons (up to 100 per carton) and provided with half the number of virgin HWE males and allowed to mate for 3-5 days. All cages were then provided with a minimum of three blood meals (defibrinated sheep blood, Colorado Serum) and allowed to lay eggs onto filter papers. G_1_ eggs were allowed to mature for five days and were then hatched, reared to fourth instar, and screened for the presence of the fluorescence marker. Positive G_1_ individuals were subsequently outcrossed to HWE again to establish the transgenic lines. Integration of the transgene was validated by PCR and Sanger sequencing using primers BR-13 and BR-115 for transgenes at the Carb109 locus and primers BR-51 and BR-73 at the TIMP-P4 locus.

#### Genetic crosses to test for GD activity

An initial pool of G_0_ hemizygous individuals was established for each of the gene drive lines by outcrossing 40 transgenic males to 200 HWE virgins *en masse* and allowing them to mate and take a blood meal. The G_0_ eggs were subsequently collected and used as the starting material for each of the drive activity crosses in our study. From the G_0_ pools, a total of 12 hemizygous males were individually crossed to two virgin HWE females, and 20-25 hemizygous females were crossed to an equal number of male HWE *en masse* and allowed to mate for 3-5 days. The individual “male founder” containers were each provided with their own blood meals, while the “female founder” container was provided with a common blood meal, followed by the transfer of blood-fed females to individual cartons for egg laying. Following oviposition, egg papers were removed and allowed to develop for a minimum of five days prior to hatching individually into 4 oz plastic cups. Larvae were reared to third/fourth instar and scored for the presence of the transgenic marker to determine the level of gene drive inheritance. Following the scoring of the outcross-1 (OX-1) mosquitoes, two pools of crosses representing either a ‘low’ level (*i.e.*, Mendelian-like), or ‘high’ level (*i.e*., Super-Mendelian-like) were selected from each cross such that an additional 12 female and 12 male G_1_ hemizygotes could be outcrossed again to HWE for the OX-2 assessment. The selection of the ‘low’ and ‘high’ pool, therefore, did not necessarily represent the lowest and highest levels of gene drive inheritance, but rather served as representative pools from which to establish the following outcross. From within OX-1, non-transgenic individuals were saved to assess for the presence of GDBI. For the AeaNosC109^GD^ and AeaZpgC109^GD^ lines, all non-transgenic mosquitoes were selected for assessment, while for the AeaNosT4^GD^ and Aeaβ2tC109^GD^ lines, a total of 20 and 10 individuals were selected, respectively. All crosses were duplicated and the data pooled for analysis.

#### Statistical analyses

Comparisons of the OX-1 gene drive levels for the lines were assessed in R using a Kruskal-Wallace non-parametric ANOVA followed by a Dunn-Bonferroni *post hoc* means separation procedure to test for significance across all groups and all lines. Comparisons of the OX-2 gene drive levels, and assessments of gene drive blocking indels were conducted in R using a Kruskal-Wallace nonparametric ANOVA followed by a Dunn-Bonferroni *post hoc* means separation procedure to test for significance among individual crosses for all groups within each line. Comparisons of the maternal and paternal contributions were assessed using pairwise T-testing in R following a Shapiro-Wilk test for normality. Deviation from the expected Mendelian inheritance level for the eCFP marker in the paternal contribution testing was conducted manually using Chi-Square analysis.

#### Testing of maternal contribution of Cas9-ribonucleo-protein (RNP)

Initial crosses to establish trans-heterozygous mosquitoes were conducted using AeaNosC109^GD^ females outcrossed to males harboring a 3xP3-eCFP-SV40 fluorescent eye marker at the Carb109 locus (**Figure S3**). The positionings of the fluorescent markers in both transgenes were in opposite orientation to one another to mitigate the potential for crossing over. Following this cross, fourth instar progeny were screened and selected for the presence of both an mCherry (gene drive) and an eCFP (null drive) marker. From this cross, 500 females of either the AeaNosC109^GD^ or the AeaZpgC109^GD^ lines were outcrossed to 200 HWE males. Approximately 1000 embryos resulting from these crosses were then microinjected with 100 ng/µL of sgRNA targeting the TIMP-P4 locus. Three rounds of injection, each > 1000 embryos were performed for each of the crosses. Embryos were allowed to develop for one week prior to hatch, then surviving larvae were reared to L3 and genotyped under a fluorescent microscope for the presence of either the eCFP marker or mCherry marker. The DNA was then individually extracted from each larva and assessed for GDBI at both the Carb109 and TIMP-P4 loci as previously described.

With this assay, any activity at the Carb109 locus in the male (non-transgenic) allele for AeaeCFPC109 mosquitoes would indicate a maternal provisioning of both the Cas9 and the sgRNA targeting the Carb109 locus. Since it was possible that the Cas9 protein could be inherited without the sgRNA, we further injected the early embryos with the sgRNA targeting the TIMP-P4 locus. This allowed for us to interrogate whether Cas9 had been supplied maternally (AeaeCFPC109 inheriting progeny) or if Cas9 had been supplied maternally and could also be expressed in the early embryo/larva (mCherry GD inheriting).

#### Testing of parental effect resulting in GD activity

To test for a parental effect that results in GD activity, trans-heterozygous crosses were performed as similarly described for the testing of the maternal effect, with the addition that crosses were performed reciprocally to include male trans-heterozygous parentals as well. The progeny from these crosses were scored for the numbers of offspring containing either mCherry or an eCFP eye marker to test for the possibility of a fitness cost at the germline level relative to the eCFP (null drive) marker. The female progeny from each of these crosses were saved (within cross) and subsequently mated to an equal number of HWE males, then provided with a single blood meal after which females were separated to allow them to lay eggs individually. Eggs were allowed to mature for at least 5 d, then hatched and scored for the presence of the eCFP marker to determine the level of transgene inheritance.

#### Assessment of GDBI

GDBI were assessed as inheritable refractory indels by analyzing the nucleotide sequence of the alternative inheritable allele in outcrossed hemizygous individuals. For the OX-1 generation of the gene drive assessment assays, non-transgenic mosquitoes were individually analyzed by PCR using primers BR-20 and BR-23 for the TIMP-P4 locus and BR-60 and BR-65 for the Carb109 locus. PCR products were amplified, purified using the Zymo Clean and Concentrator kit (Zymo Research, San Diego, CA), and Sanger sequenced. Chromatogram traces were then assessed manually for trace decay at the predicted CRISPR/Cas9 cleavage site. For the assessment of GDBI in the OX-2 generation, all non-transgenic mosquitoes within a cross were pooled and their DNA extracted to obtain an ‘average’ number of GDBI for the non-drive-inheriting mosquitoes. PCR products were then amplified using primers BR-724 and BR-725 for the Carb109 locus and BR-726 and BR-727 for the TIMP-P4 locus to obtain the ‘round 1’ PCR products for Illumina whole-amplicon sequencing. The first round PCR was held to 30 cycles. PCR products were subsequently purified using the Zymo Gel Extraction kit (Zymo Research, San Diego, CA) and adjusted to 1 ng/uL using a Qubit fluorimeter (ThermoFisher Scientific, Waltham MA), and material was subsequently sequenced using paired end reads of 250 bp length (PE250) on an Illumina MiSeq instrument. The resulting reads were trimmed using Cutadapt v1.01 (39) and assessed for indels using Crispresso v2 (40). The output files of Crispresso v2 containing insertions, deletions, and substitutions (assumed to be a composite event of deletion/insertion) were then summed up to generate the value for ‘indels’.

### Fitness cost studies

Life parameter data was collected from male hemizygote AeaNosC109^GD^ and AeaZpgC109^GD^ parentals as well as from HWE for comparison. Each fitness measurement was calculated as an average from a minimum of 100 individual egg papers per mosquito strain ± standard error of the mean. All data were analyzed with one-way ANOVA, and if a significant difference was found, one-tailed t-tests were performed comparing HWE and each transgenic line. All mosquitoes were maintained at 28 °C with 75 –80% relative humidity and a 12 h light / 12 h dark cycle. Data are displayed in **Table S7**.

#### Fecundity

Three to five days post-emergence, hemizygous males or HWE males were mated *en masse* with virgin HWE females in ratios of 5 males : 1 female using 64 oz. cartons (WebstaurantStore, catalog number: 76 999SOUP64WB) each containing ∼ 150 mosquitoes. Around 2-3 days later, females were offered an artificial blood meal containing defibrinated sheep blood (Colorado Serum Co., Denver, CO, USA) and 10 mM ATP, after which engorged females were retained. Two to three days later, individual females were placed in 50 ml conical tubes lined with Whatman Grade 1 Qualitative Filter Paper (catalog number: 1001-824) and filled with ∼ 5 ml tap water. Females were allowed to lay eggs for 1-2 days. Papers were visually inspected for eggs, which were then quantified for fecundity, defined as the number of eggs a female oviposited in one gonotrophic cycle. Egg papers were dried for a minimum of 5 days.

#### Fertility, sex ratios, larva viability, and larva-to-pupa development

Individual egg papers were hatched in ∼ 100 ml freshly sterilized deionized water using ProPak 1600 polypropylene clear deli containers (Webstaurant) and were offered a ground fish food (Tetramin, Melle, Germany) slurry *ad libitum*. Around 2-3 days post-initial hatch, the egg papers were removed from the water and were allowed to dry again overnight. All egg papers were then rehatched in the same containers from which they were initially hatched. Larvae (2^nd^-4^th^ instars) were visually inspected under a fluorescent microscope for expression of the mCherry eye marker. Both positive and negative larvae were counted, but negative larvae were eventually discarded. Fertility was calculated as the total number of larvae / total number of eggs. Female and male pupae were collected every day, up to 14 days post-hatch, and the timepoints of pupation were recorded. Around 150 mosquitoes (either males or females) of each mosquito line were each pooled into small cups and allowed to emerge in 64 oz cartons. Sex ratio was defined as the total number of males or females / the total number of pupae; larva viability was defined as the total number of pupae / total number of eggs; and larva-to-pupa development was defined as the average time to pupae development post-egg hatch.

#### Male competition and adult longevity

Using virgin adults, 50 transgenic and 50 HWE males (age matched) were allowed to mate with 100 HWE females for 2-3 days. Females were then offered an artificial blood meal containing defibrinated sheep blood and 10 mM ATP. Eggs from individual engorged females were obtained as described above for the fecundity assay. Around 5 days later, the egg papers were hatched in sterile water, and larvae were fed ground Tetramin fish food *ad libitum*. Larvae (2^nd^-4^th^ instars) were visually inspected under a fluorescent microscope for the expression of the mCherry eye marker. Male competition data was calculated as the number of positive G_2_ larvae / total number of larvae. For the longevity assay, 50 male and 50 female transgenic mosquitoes (3-5 days post-emergence) were maintained in 64 oz. containers and offered raisins (as a sugar source) and water. Every day, the number of dead mosquitoes was recorded and longevity was calculated as the average number of days passed when 50% of the male or female mosquitoes had died across cartons.

### MGDrivE population modeling

The mosquito GD explorer (MGDrivE v 1.6.0) was used to model the inheritance of the AeaNosC109^GD^ and AeaZpgC109^GD^ lines (41). A stable population was set to 10000 mosquitoes at a 1:1 sex ratio with no migration out of, or into, the main patch. A single release of 2000 homozygous male mosquitoes (20%) was simulated at day 25 for 100 stochastic samplings for 1500 days (AeaNosC109^GD^) and 4000 days (AeaZpgC109^GD^). In the MGDrivE package, the cube “Cube-CRISPR2MF.R” allows for sex-specific rates of drive and allows for the modeling of two types of resistance alleles: those that block the GD, yet cause no fitness cost (R), and those that block the GD but result in a fitness cost (B). Since it was predicted that a GDBI occurring in an intergenic locus would cause no fitness cost, we modified the tMatrices in the cube script “Cube-CRISPR2MF.R” to still allow for sex-specific rates of drive and indel causing activities, but we condensed the (R) and (B) alleles to a common (R) allele, which represented all GDBI and assumed no fitness cost to that genotype. The modified gtype and tMatrices are provided as **Table S8**. We further plotted an additional figure panel that summed females with any combination of genotypes containing a GD allele and any combination of genotypes without a GD allele to simulation. These values were used to estimate the proportion of a hypothetical linked antiviral effector gene to represent females having a viral refractory phenotype. The parameters used for our model and how they were calculated are provided in **Table S9**. The entomological parameters popGrowth and muAD were left as default, based on literature. tEgg and tPupa were assumed to be 5 and 2, respectively, based on standard laboratory rearing practices and observations. tLarva was calculated as the average number of days of larva-to-pupa development. betaK was calculated as the average fecundity * 4 (the average number of blood meals a female takes during her lifetime) / 11 (the average lifespan of a female mosquito in an urban environment) (42). They were determined from Table 3 by adding the marker positive mosquitoes from the “low” and “high” pools together to obtain the total number of correct homing events. The number of incorrect homing events (*i.e*., GDBI formation) was estimated by multiplying the total number of marker-negative mosquitoes from the “low” and “high” pools by the fraction of the tested OX-1 marker-negative progeny identified to contain GDBI. This was done in place of a direct count since some of the samples from the OX-1 marker-negative larvae did not provide a PCR product that could be sequenced for the assessment of GDBIs. Then, to calculate the parameters cF and cM, the number of marker positive and GDBI positive mosquitoes were summed and divided by the total N. From this number, the proportion that gave rise to correct homing events was determined as chF and chM, with the resistance rate formation parameters crF and crM calculated as 1-chF or 1-chM, respectively. The rate of maternal deposition, dF, was calculated as the average rate of GDBI observed for the Carb109 locus for eCFP-inheriting offspring from the trans-heterozygous cross between the GD and the eCFP-blocked GD. For both the AeaNosC109^GD^ and AeaZpgC109^GD^ line, we then further set the rate of correct maternal effect homing (dhF) to 0 and the rate of incorrect maternal effect homing (drF) to 1. Larval viability was significantly reduced for the AeaNosC109^GD^ line, which we incorporated into the model under the xiF/xiM variables by subtracting the average larval viability of the HWE strain from the average larval viability of the AeaNosC109^GD^ line. In the AeaZpgC109^GD^ line, the larval viability was slightly higher than in HWE, but not significantly different. Therefore, we set the parameters xiF and xiM to NULL for the AeaZpgC109^GD^ line. Finally, since Li *et al*. (2020) similarly found that their transgenic split GD had an overall fitness similar to the parental Liverpool strain, yet still included a 10% fitness cost for their modeling, we incorporated an overall decrease in fertility by 10% for both of our modeled GD lines (10).

## RESULTS

### Identification of active target sites and target site polymorphisms

Prior to testing the GD lines, we first identified two targets for CRISPR/Cas9 cleavage, which were located in non-protein encoding regions of the mosquito genome. In our previous work (17), we identified a highly active sgRNA for the TIMP-P4 locus (Table 1). In our current work, we identified a novel active CRISPR/Cas9 target site for the Carb109 locus located 1214 bp downstream of the *mariner/Mos1* insertion site, which is the defining locus in the DENV2-resistant Carb109 line (13). Since the Carb109 *mariner/Mos1* insertion was determined to be within the 3’-UTR of the polyadenylate-binding protein gene (AAEL010318), we manually selected CRISPR/Cas9 targets as close to, but not within, the AAEL010318 gene locus and assessed for off-targets using CHOPCHOP v2 (43). In total, we assessed six sgRNA for activity (Table 1) and identified only a single sgRNA (#15) that showed DNA cleavage activity. Therefore, for all CRISPR/Cas9 based experiments at the Carb109 locus, sgRNA #15 was selected. While sgRNA #15 was ideal for our GD testing in the laboratory-adapted HWE strain of *Ae. aegypti*, we further wanted to estimate how conserved the sgRNA #15 target was in field strains of this mosquito species. In addition to the assessment of sgRNA activity, we therefore also analyzed both genomic target sites, TIMP-P4 and Carb109, for polymorphisms using data from Schmidt *et al* (2020), which examined CRISPR/Cas9 target sites across 132 genomes of *Ae. aegypti* (44). Two polymorphisms were found in the TIMP-P4 locus: one in the protospacer adjacent motif (PAM) that changed the second base G to a T and had an allele frequency of 0.026 among the 132 genomes tested, while the second polymorphism was at position 13 distal to the PAM that changed the base from a G to a C at an allelic frequency of 0.009 (**Table S1**). The Carb109 locus contained only a single polymorphism at position 19 distal to the PAM that still allows for the use of an 18 bp protospacer, which is considered functional for Cas9 activity (45). Although most of the data from Schmidt et al (2020) is derived from *Ae. aegypti* populations in California, we assumed no polymorphisms for the TIMP-P4 and Carb109 loci in regard to our population modeling.

**Table 1.** Active CRISPR/Cas9 target sites proximal to the original TIMP-P4 and Carb109 *mariner/Mos1* insertion loci.

| Locus | sgRNA number | Genomic locus (distance from original *mariner/Mos1* insertion site) | Activity (# embryo pools / total pools) |
| --- | --- | --- | --- |
| TIMP-P4 | sgRNA #5* | 2: 321382225 (623 bp) | Active (3/3) |
| Carb109 | 2 | 3: 409717866 (19942 bp) | Not active (0/3) |
|  | 4 | 3: 409723215 (25291 bp) | Not active (0/3) |
|  | 6 | 3: 409699241 (1317 bp) | Not active (0/3) |
|  | 15 | 3: 409699138 (1214 bp) | Active (3/3) |
|  | 21 | 3: 409699885 (1961 bp) | Not active (0/3) |
|  | 29 | 3: 409699526 (1602 bp) | Not active (0/3) |

\*Previously reported in Williams *et al.*, 2020 (17)

### Establishment of transgenic Ae. aegypti lines

A total of six transgenic lines were established via homologous recombination with embryo injection efforts ranging from 568 – 1442 embryos (Table 2). In general, the survival rate of the injected G_0_ mosquitoes ranged from a low of 3.9% (AeaeCFPC109) to a high of 23% (AeaNosC109^GD^). In addition, we found that the *nanos*Cas9-U6sgRNA constructs required no helper Cas9 or helper sgRNA to integrate into the genome. Due to the timing of expression of native *nanos*, there was co-expression of Cas9 from the injected plasmid, and due to the ubiquitous expression of the AAEL017774 U6 gene, there was co-expression of sgRNA as well. Regardless, after the initial successful injection of the *nanos*Cas9-U6sgRNA construct into the TIMP-P4 locus, helper Cas9 protein was added to all injection mixes, with the omission of the helper sgRNA and continued inclusion of the ku70 dsRNA trigger.

**Table 2.** *Aedes aegypti* lines used in our study and respective injection efforts for their establishment.

| Line name | Marker | Purpose | Embryos injected | Survivors (F:M ; %) |
| --- | --- | --- | --- | --- |
| Aeaβ2tT4<sup>GD</sup> | 3xP3 mCherry | GD at TIMP-P4 locus | 1115 | 72 (31:41 ; 6.5%) |
| AeaNosT4<sup>GD</sup> | 3xP3 mCherry | GD at TIMP-P4 locus | 959 | 154 (66:88 ; 16%) |
| Aeaβ2tC109<sup>GD</sup> | 3xP3 mCherry | GD at Carb109 locus | 882 | 122 (64:58 ; 13%) |
| AeaNosC109<sup>GD</sup> | 3xP3 mCherry | GD at Carb109 locus | 912 | 206 (90:116 ; 23%) |
| AeaZpgC109<sup>GD</sup> | 3xP3 mCherry | GD at Carb109 locus | 568 | 45 (17:28 ; 7.9%) |
| AeaeCFPC109 | 3xP3 eCFP | Genetic balancer for Carb109 locus | 1142 | 44 (24:20 ; 3.9%) |

### GD inheritance rates over two outcrossed generations

To test for the rates of GD inheritance, a starting population was established by outcrossing transgenic males to non-transgenic HWE females to establish hemizygous “OX-0” (outcross 0) mosquitoes. This was performed to ensure that all mosquitoes used for testing were hemizygous. Overall, the strongest GD performance was observed in the AeaNosC109^GD^ line, where the median inheritance rate among progeny was 70% and 73% for male and female parentals, respectively (Figure 1; **Table S2**). Further, the levels of drive ranged from no drive, 45-48%, (comparable to Mendelian inheritance rates) to super-Mendelian rates, reaching up to 96% (Figure 1; **Table S2**). When the same cargo was inserted and retargeted for the TIMP-P4 locus, however, no GD activity was observed with median values for marker inheritance remaining at 48% and 46% for male and female parentals, respectively (Figure 1; **Table S2**). Taken together, these results show that *nanos*Cas9 GDs were principally functional in *Ae. aegypti* and overall GD activity was strongly influenced by the GD’s genomic insertion site. Regarding the *β2tubulin*Cas9 controlled GD construct, the median values for marker inheritance demonstrated no drive activity for the Aeaβ2tC109^GD^ line, with overall transgene inheritance rates resembling Mendelian inheritance patterns as they ranged from a low of 43% in male parentals (n = 24/56) to a high of 55% in both male (n = 16/29) and female (n = 11/20) parental groups. Given that GD activity was observed at the Carb109 locus using the *nanos* promoter, we tested another promoter in the same locus, *zero population growth (zpg)*, which has been shown to have strong GD activity in *An. gambiae* but has not yet been tested in *Ae. aegypti* (46, 47). The median GD inheritance rates for the AeaZpgC109^GD^ line were 56% and 68% for male and female parentals, respectively (Figure 1; **Table S2**). The male transgenic parentals had a significantly lower (*p* = 0.0258) level of drive when compared to either parental cross for the AeaNosC109^GD^ line, or to female transgenic parentals of the AeaZpgC109^GD^ line. These results showed that while the *zpg* promoter is active in *Ae. aegypti*, there may be a sex-specific effect on Cas9 expression, which was not anticipated for this promoter since the *zpg* gene (*innexin-4*) is involved in early germ cell development in both male and female *Drosophila* (48). Further, given that the median GD inheritance rates for the Aeaβ2tC109^GD^ line were below 50% for both male and female parentals, it suggested that there was no CRISPR/Cas9 activity in the germline for this line. Recent work by Terradas *et al* (2021) indicated that in *An. gambiae*, the *β2-tubulin* promoter is expressed post-meiotically, which could explain why we did not see GD activity when using this promoter (49). Therefore, we did not continue the Aeaβ2tC109^GD^ line for further assessment at the OX-2 level. Meanwhile, although the AeaNosT4^GD^ line also did not demonstrate any GD activity, its cargo was identical to that of the AeaNosC109^GD^ line, prompting us to continue this line for further assessment at the OX-2 level.

**Figure 1.**
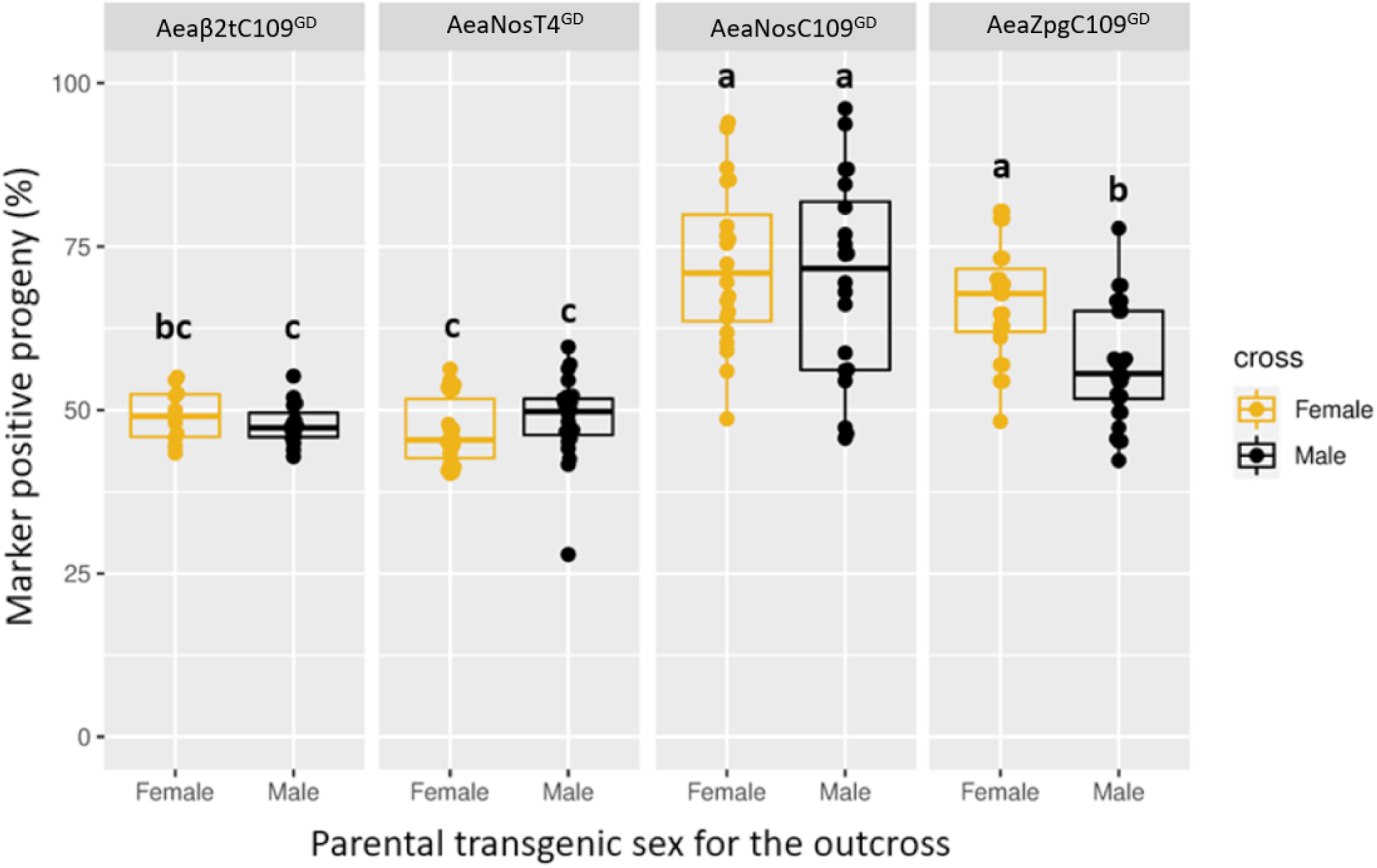
Frequency of marker gene inheritance in first-outcrossed (OX-1) individuals originating from hemizygous outcrossed females or males of transgenic lines harboring CRISPR/Cas9 based GDs at two different genomic loci (Carb109 and TIMP-P4) with Cas9 expression under control of three different promoters (β*2tubulin*, *nanos*, or *zpg*). The bar inside each box plot represents the median value, while the lower and upper borders of each box represent Q1 and Q3, respectively. Groups superseded with the same letter are not statistically different (*p* > 0.05). Each data point represents the percentage of transgene inheritance resulting from the offspring of the parental crosses where each transgenic female parental was allowed to mate with two non-transgenic males, and each transgenic male parental was allowed to mate with two non-transgenic females. A minimum of 20 larvae was set for each group in order to be scored. For the female parental crosses, the population sizes of each data point ranged from 20 to 142 larvae (mean ± SEM = 59 ± 3 larvae). For the male parental crosses, the population sizes of each data point ranged from 21 to 415 larvae (mean ± SEM = 91 ± 5 larvae).

Regarding the OX-2 crosses, we were interested in answering two questions: 1) Would the effects of ‘low’ and ‘high’ levels of GD inheritance from the OX-1 crosses be consistent? and 2) Do GD blocking indels (GDBI) accrue differently for female and male transgenic parentals? We assessed for these by first identifying individual crosses from each of the OX-1 groups that represented ‘low’ and ‘high’ levels of drive (Table 3), and subsequently outcrossed transgenic female and males reciprocally to HWE mosquitoes as was performed in OX-1. This resulted in a total of eight crossing groups within each line to which we applied three-letter codes that identified the parental (F ‘female’ or M ‘male’), the grandparental (F ‘female’ or M ‘male’), and the level of drive observed for the OX-1 pool (L ‘low’ or H ‘high’). When the progeny (n ranging from 795 to 3162) for all OX-2 crosses were scored, we again observed Mendelian-like inheritance of the GD cassette for the AeaNosT4^GD^ line. However, one cross in the FFH group resulted in 90% inheritance (n = 124), suggesting that while GD activity was inefficient at the TIMP-P4 locus, occasional cases of drive might still occur (Figure 2; **Table S3**). The results for the AeaZpgC109^GD^ line displayed similar levels of drive for all 157 crosses, with median inheritance rates ranging from 50% in the MML group to 60% in the FMH group (Figure 2; **Table S3**). Further, among all AeaZpgC109^GD^ crosses, five groups contained individual crosses that demonstrated > 80% rate of GD inheritance (Figure 2; **Table S3**). These results suggest that the effect of ‘low’ or ‘high’ level of drive was not inheritable by the OX-2 generation. Finally, for the AeaNosC109^GD^ line, the median GD inheritance rates ranged from a low of 56% in the FML group to a high of 77% in the MFL group (Figure 2; **Table S3**). Of particular interest was that in the AeaNosC109^GD^ line, there were outcrosses that occasionally resulted in 100% inheritance of the GD. While these events were rare, they were found across multiple parental and grandparental groups, including the MMH, MFH, FFL, and MFL groups (n = 20, 44, 114, and 137, respectively) (Figure 2; **Table S3**). These results highlight that while, overall, the crossing groups within line AeaNosC109^GD^ line displayed GD inheritance rates ranging from 57% to 76% (**Table S3**), complete drive (100% inheritance) was achievable. Given that the levels of GD in the AeaNosC109^GD^ line were statistically similar (*p* > 0.05) for male parentals with any grandparental combinations irrespective of their ‘low’ or ‘high’ level of GD origins, we ruled out the effect of grandparent sex and the possibility that the trait of ‘low’ or ‘high’ GD levels was inheritable.

**Figure 2.**
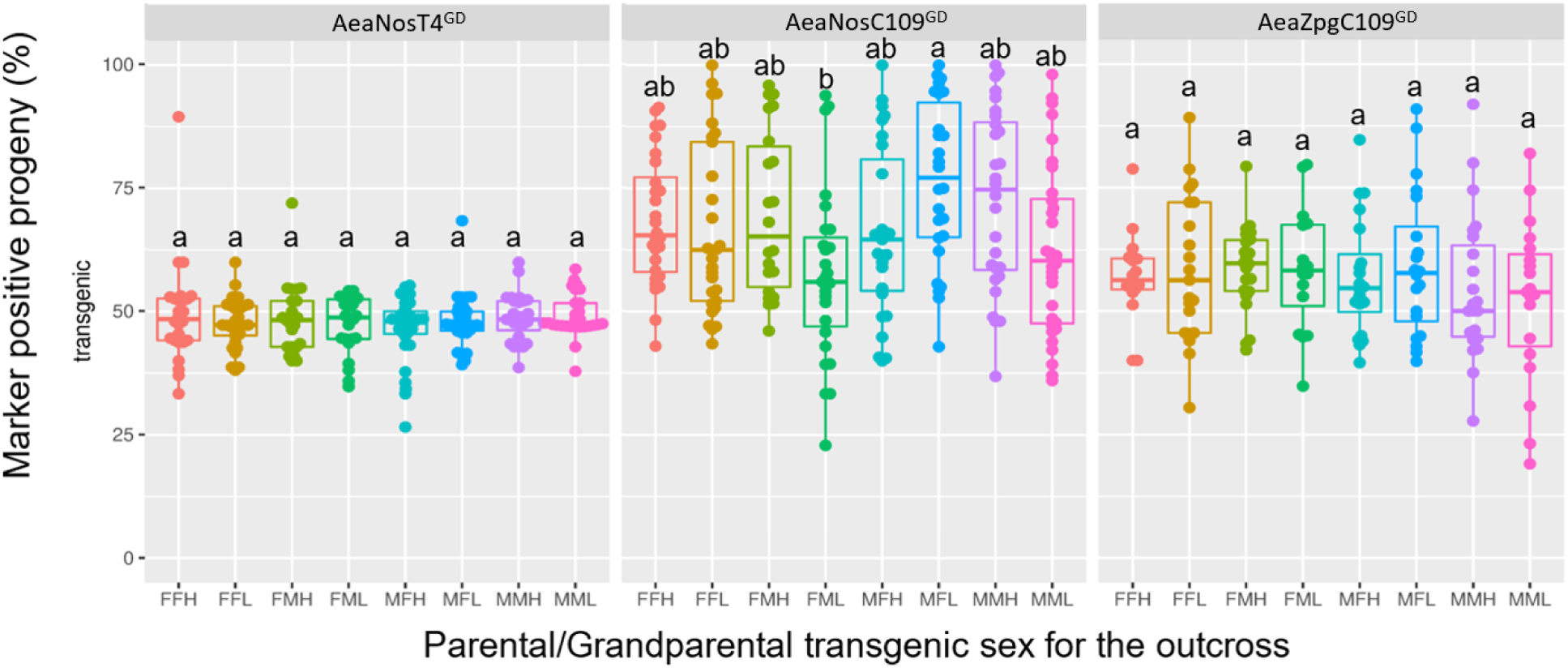
Frequency of marker gene inheritance in progeny from second-outcrossed (OX-2) individuals (594 groups, average n = 72 ± 2, total n = 43,015) originating from either ‘high’ or ‘low’ parental GD pools harboring the GD at two different genomic loci (Carb109 and TIMP-P4) with Cas9 expression under control of two different promoters (*nanos* or *zpg*). The bar inside each box plot represents the median value, while the lower and upper borders of the boxes represent Q1 and Q3, respectively. Groups within a GD line that are superseded with the same letter are not statistically different (*p* > 0.05). The first letter of each cross designation indicates the OX-2 parental transgenic sex and the second letter indicates the OX-1 grandparental transgenic sex, where F=female and M=male. L=Low/no drive level in the grandparental generation, H=high drive level in the grandparental generation. Each data point represents the percentage of transgene inheritance resulting from the offspring of the parental crosses where each transgenic female parental was allowed to mate with two non-transgenic males, and each transgenic male parental was allowed to mate with two non-transgenic females. A minimum of 20 larvae was set for each group in order to be scored.

**Table 3.** Testing of non-transgenic offspring from OX-1 individuals to assess for the presence of GD blocking indels (GDBI) among the GD lines.

| Line | GD Parental | GD inheritance rate<br>(% marker inheritance; n<br>total) | Number of GD-negative<br>larvae with indels / total<br>number of GD negative larvae<br>assessed (%) |
| --- | --- | --- | --- |
| AeaNosT4<sup>GD</sup> | Male | Low (45.9%; 290) | 0/40 (0%) |
|  |  | High (58.1%; 136) | 0/40 (0) |
|  | Female | Low (42.9%; 163) | 0/40 (0) |
|  |  | High (51.3%; 117) | 0/40 (0) |
| AeaNosC109<sup>GD</sup> | Male | Low (55.3%; 228) | 9/54 (16.7) |
|  |  | High (86.4%; 214) | 7/27 (25.9) |
|  | Female | Low (61.5%; 148) | 7/46 (15.2) |
|  |  | High (89.7%; 136) | 7/12 (58.3) |
| AeaZpgC109<sup>GD</sup> | Male | Low (56.5%; 232) | 0/41 (0) |
|  |  | High (59.2%; 228) | 0/51 (0) |
|  | Female | Low (57.1%; 98) | 2/27 (7.4) |
|  |  | High (73.7%; 118) | 1/23 (4.3) |
| Aeaβ2tC109<sup>GD</sup> | Male | no GD (50.7%; 138) | 0/10 (0) |
|  | Female | no GD (52.5%; 40) | 0/10 (0) |

### Accrual of GD blocking indels (GDBI)

To better understand why a broad range in GD activity was observed, we assessed for the occurrence of GDBI among the OX-1 and OX-2 crosses. Overall, the occurrence of GDBI in the OX-1 crosses was most pronounced in the AeaNosC109^GD^ line, which also demonstrated the highest average level of GD (Figures 1,2; Table 3). Meanwhile, the *zpg* promoter similarly displayed some GDBI, albeit at a much lower level (7.4% for ‘low’ GD individuals, and 4.3% for ‘high’ GD individuals), and only in pools of offspring originating from females. For both the Aeaβ2tC109^GD^ and AeaNosT4^GD^ lines, no GDBI were identified in the OX-1 crosses; however, these lines did not demonstrate GD activity and had Mendelian-like inheritance of the marker; thus, it was not expected that GDBI would arise in these lines as there was likely no CRISPR/Cas9 activity. Given that the GDBI identified in the OX-1 crosses were inherited from the original transgenic males, which had been crossed to HWE females in the OX-0 cross, we further assessed the levels of GDBI for the OX-2 crosses to identify any differences between female and male groups with respect to the number of GDBI that arose in the population (Figure 3; **Table S4**). Overall, we identified negligible proportions (< 5%) of amplicons containing GDBI in the AeaNosT4^GD^ line, comparable to the single HWE control sample (Figure 3). This observation was in accordance with the results of the OX-1 GDBI assessment, and again was likely due to the lack of activity of CRISPR/Cas9 when the GD was positioned at the TIMP-P4 locus. Regarding the assessment of the OX-2 GDBI in the AeaNosC109^GD^ line, all crossing groups displayed variable proportions of amplicons containing GDBIs. In addition, we observed nine crosses for which the proportion of amplicons containing GDBI was 50%. A proportion of 50% represents the theoretical maximal proportion of amplicons that can contain indels when considering outcrossed (GD-negative) larvae. Of these nine crosses, two data points were close to 50% (FFL: 54%, MFL: 53%), while the remaining seven data points ranged from 64% (MMH) to 100% (MML). These elevated proportions of amplicons containing r GDBI could be an artifact of PCR chimera. Thus, our analysis of pooled larvae based on high-throughput sequencing was likely overestimating the actual number of GDBI in the sample. One crossing group within the AeaNosC109^GD^ line, MFH, contained very few amplicons showing the presence of indels (median = 14.5%); however, the median proportion of amplicons containing indels was not significantly lower in this group than in the other crossing groups within the AeaNosC109^GD^ line (*p* = 0.0831). Finally, in the AeaZpgC109^GD^ line, a significantly greater median number of GDBI was observed for the FFL group (median = 2.6%) when compared to the MMH (median = 0.02%) group. In general, all male parental crosses appeared to have fewer GDBI (median range = 0.02% - 1.6%) than the female parental crosses (median range = 2.0% - 11.9%), regardless of the sex of the grandparental transgenic individual. As explained above, this result was unexpected for the *zpg* promoter (48). Interestingly, there was no apparent grandparental effect for the *zpg*Cas9 controlled GD. In addition to the abundance of GDBI assessed for the OX-2 cross, we also assessed the size distributions of the resultant insertions/deletions (**Figure S1**). Overall, deletions were the most abundant type of indel, representing 86%, 94%, and 97% of all indels for the AeaNosT4^GD^, AeaNosC109^GD^, and AeaZpgC109^GD^ lines, respectively. Nearly all identified deletions were present within 50 bp of the CRISPR/Cas9 target site, with the largest deletion being 82 bp in the AeaZpgC109^GD^ line. The largest identified insertion was 60 bp in the AeaNosC109^GD^ line, although nearly 100% of the insertions in all lines tested were within 50 bp of the CRISPR/Cas9 target site. Taken together, these results suggest that for the purpose of an autonomous GD system in *Ae. aegypti*, the homologous region required for strand invasion of the cleaved DNA proximal to the predicted CRISPR/Cas9 target is likely a minimum of 50 bp in size.

**Figure 3.**
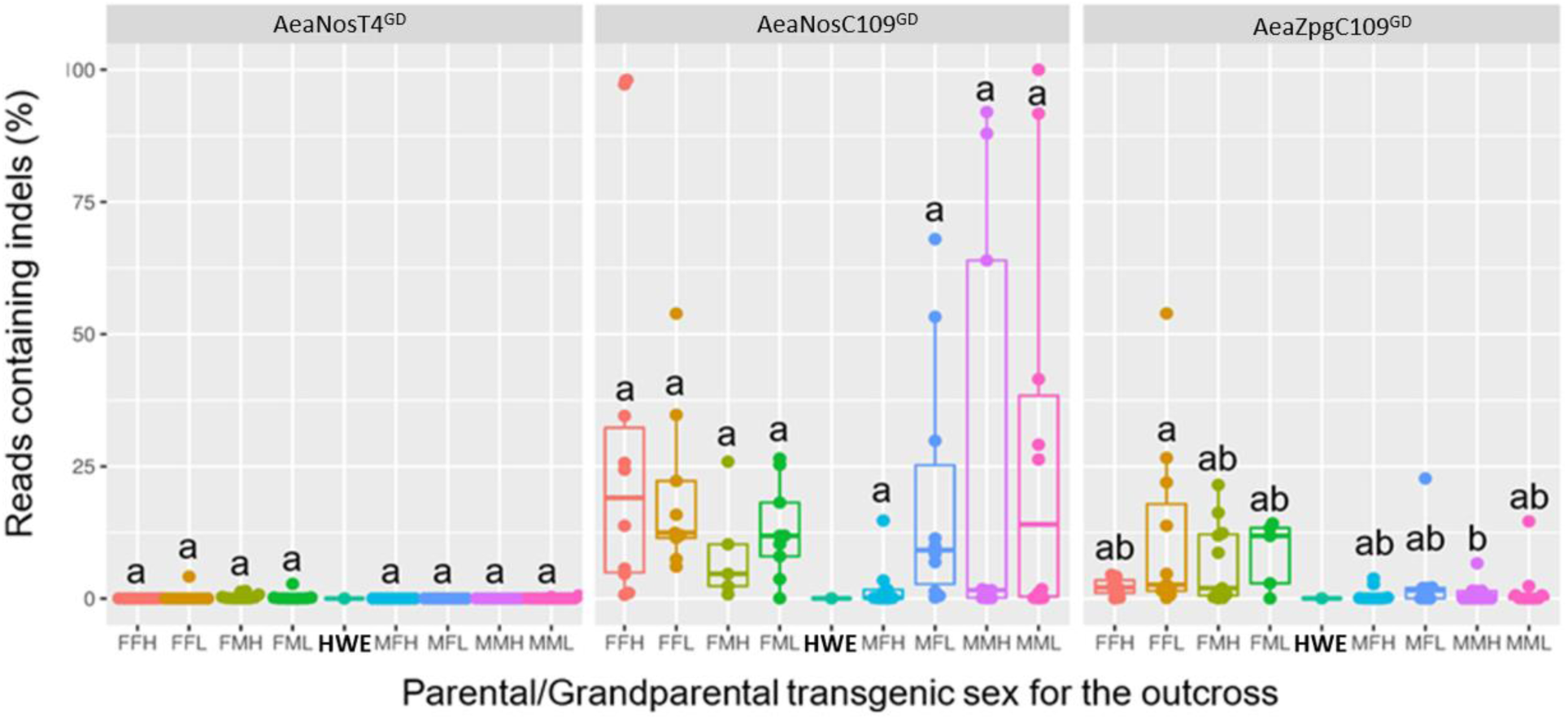
Proportions of PCR amplicons containing GDBI for pooled non-transgenic mosquitoes from the second-outcrossed (OX-2) GD lines originating from either ‘high’ or ‘low’ parental GD pools harboring the GD at two different genomic loci (Carb109 or TIMP-P4) and Cas9 expression under control of two different promoters (*nanos* or *zpg*). The bar inside the box plot represents the median value, while the lower and upper borders of the boxes represent Q1 and Q3, respectively. Indels were analyzed via high-throughput nucleotide sequencing. The first letter within each cross indicates the parental transgenic (OX-2) sex, the second letter indicates the grandparental (OX-1) transgenic sex. L=Low/no drive level in the parental generation, H=High drive in the parental generation. HWE=Higgs’ white eye non-transgenic. Each data point represents the percentage of high-throughput reads containing a GDBI obtained from PCR amplicons that spanned the genomic target site for either Carb109 or TIMP-P4 locus. The gDNA template used for each PCR reaction consisted of the pooled contribution of all non-transgenic larvae from the OX-2 crosses.

### Testing for maternal contribution of CRISPR/Cas9

Maternal and/or paternal contribution of the CRISPR/Cas9 components leads to extra-germline activity of the Cas9 ribonucleo-protein (RNP) in the early embryo, which could result in the development of GDBI. Notably, this will occur in offspring that do not inherit the GD allele, which in the absence of a donor for HDR would cause GDBI formation. To test for maternal contribution, the paternal allele must be distinctly identifiable from the inherited maternal allele when the GD allele is absent. Previous studies have used recessive phenotypes to identify these effects (10, 50–51), where the maternal allele is balanced over a null-allele for the recessive marker and the female is then outcrossed to a wildtype male. The progeny from this cross is phenotypically wildtype and genotypically heterozygous. In the event of cleavage of the paternal allele, however, the progeny will be biallelic/homozygous and display the recessive phenotype. Since we targeted an intergenic locus, we accomplished the same goal through the development of the AeaeCFPC109 line, which contained a marker-only cargo and no GD construct: a codominant alternate color marker (eCFP). This construct was then integrated into the *Ae. aegypti* genome through homologous recombination using sgRNA #15 and was therefore positioned at the same (Carb109) locus as the mCherry-marked GD systems. Since the sgRNA #15 target site was interrupted by the 3xP3-eCFP marker, it established a “blocked drive” because the sgRNA #15 protospacer is no longer present in the genome. When the AeaeCFPC109 line is crossed to either the AeaNosC109^GD^ or the AeaZpgC109^GD^ lines, offspring containing both eCFP and mCherry eye markers would harbor one allele with an eCFP marker, and the other allele with the mCherry-marked GD. No drive or GDBI activity would occur in the germline as the genomic GD target is blocked by the eCFP marker meaning that the Carb109/sgRNA #15 locus is no longer present (**Figure S3**). The genotypes of the eggs of trans-heterozygous females resulting from the cross will then consist of either the mCherry-marked GD or the GD blocking eCFP marker. When these trans-heterozygous females are outcrossed to non-transgenic male HWE mosquitoes, the males will contribute an allele that is targetable by CRISPR/Cas9 in the embryo. This then can allow for GD/cleavage activity to be assessed. Given that CRISPR/Cas9 activity requires both the Cas9 protein and the sgRNA transcript, we further microinjected embryos from the cross with synthetic sgRNA targeting a secondary genomic locus (TIMP-P4 locus). These assays allowed us to reveal any maternal effect in the early embryo and this way provided insight to possible early embryo (extra-germline) activity of CRISPR/Cas9 components. Three sets of approximately 1000 embryos were injected for each of the HWE outcrossed trans-heterozygotes (WT X [AeaNosC109^GD^/ AeaeCFPC109], and WT X [AeaZpgC109^GD^ /AeaeCFPC109]) and allowed to develop to third instar to enable visual genotyping of the fluorescent marker (Table 4). DNA was extracted from individually surviving mosquitoes and PCR followed by Sanger sequencing was conducted across both the Carb109 and TIMP-P4 loci to assess for the presence of GDBI at the target loci. We observed for both the AeaNosC109^GD^ and the AeaZpgC109^GD^ lines CRISPR/Cas9 activity in the GD-targeted Carb109 locus and in the TIMP-P4 locus targeted via injected sgRNA. We also found that the survival rate of sgRNA-injected trans-heterozygotes was lower when compared to that of the sgRNA injected (non-transgenic) HWE strain, which could be due to the concurrent targeting of two loci on different chromosomes.

**Table 4.** Embryos injected for maternal contribution testing and the resultant genotype frequencies of surviving larvae used for CRISPR/Cas9 activity assessment.

| Cross | Rep | Embryos injected | Surviving larvae N, (% of injected) | eCFP N, (% of larvae) | mCherry N, (% of larvae) |
| --- | --- | --- | --- | --- | --- |
| [AeaNosC109<sup>GD</sup> x | 1 | 1056 | 26 (2.4) | 14 (53.8) | 12 (46.1) |
| AeaeCFPC109] | 2 | 1017 | 33 (3.2) | 18 (54.5) | 15 (45.5) |
|  | 3 | 1170 | 24 (2.0) | 14 (58.3) | 10 (41.7) |
|  | 1 | 1007 | 23 (2.3) | 13 (56.5) | 10 (43.5) |
| [AeaZpgC109<sup>GD</sup> x | 2 | 1015 | 26 (2.6) | 12 (46.1) | 14 (53.8) |
| AeaeCFPC109] | 3 | 1004 | 46 (4.6) | 22 (47.8) | 24 (52.2) |

In general, the CRISPR/Cas9 activity observed for maternal deposition (inheriting the eCFP eye marker) was highest in the AeaNosC109^GD^ line, with 40% of the larvae demonstrating Cas9 activity at the TIMP-P4 locus and 19% displaying Cas9 activity at the Carb109 GD locus (Figure 4). In the AeaZpgC109^GD^ line, 16% and 13% of larvae exhibited Cas9 activity at the TIMP-P4 and Carb109 sites, respectively, although statistically, these values were not significantly different across genomic targets or mosquito lines (*p* > 0.05) (Figure 4). We did, however, find that in the AeaNosC109^GD^ line, significantly more indels were detectable at the Carb109 locus when the GD allele was inherited (*p* = 0.0309). In addition, there were significantly more indels present among GD inheriting progeny from the AeaNosC109^GD^ line than among GD inheriting progeny arising from the AeaZpgC109^GD^ line crosses (*p* = 0.0051). Taken together, these results suggest a clear and strong maternal inheritance of the CRISPR/Cas9 RNP. Furthermore, early embryonic activity of Cas9 was stronger when additional sgRNA was supplied, either exogenously through microinjection of synthetic TIMP-P4 targeting sgRNA, or endogenously as Pol-III expressed sgRNA in the GD inheriting larvae.

**Figure 4.**
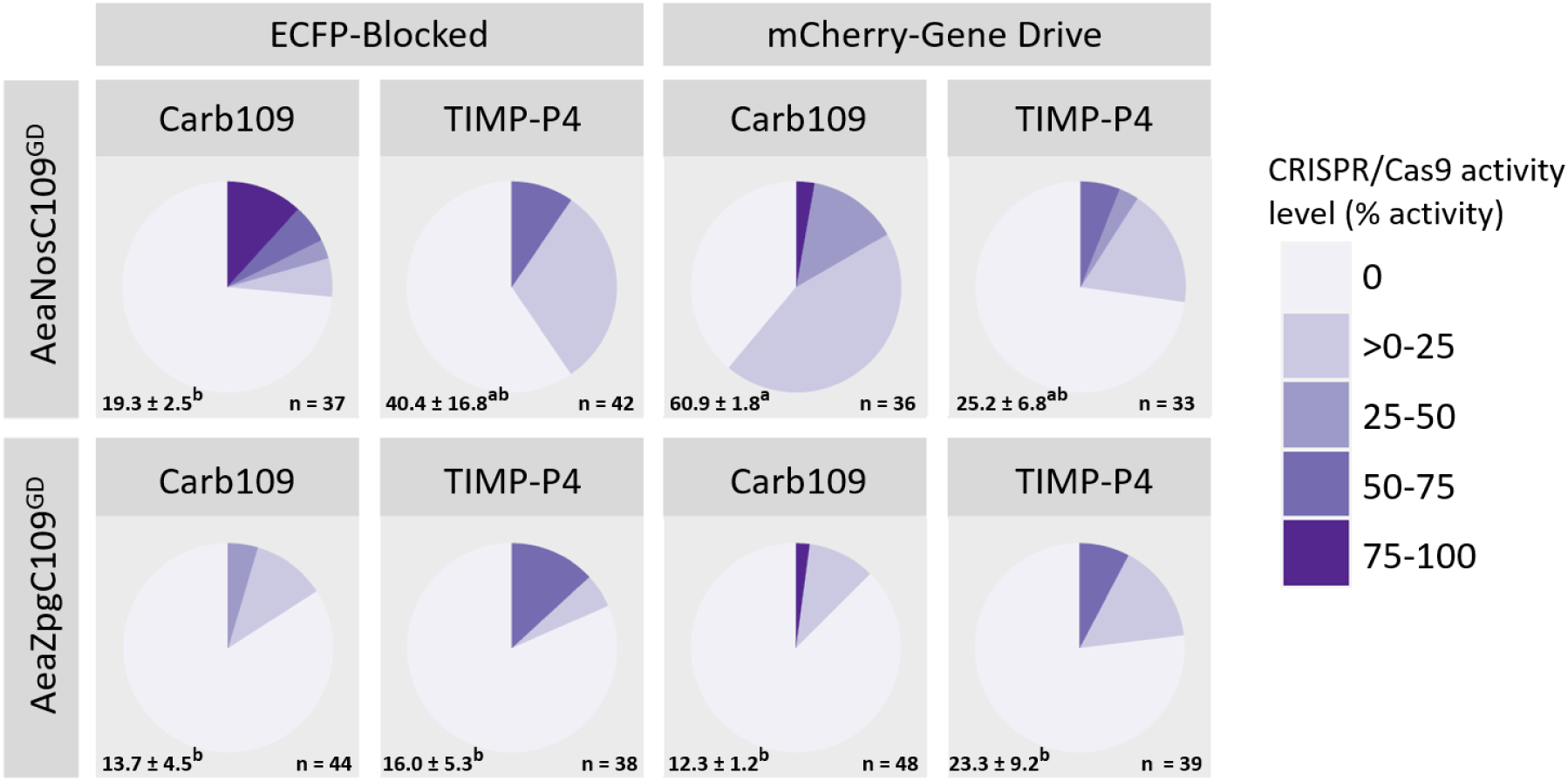
Maternal contributions of CRISPR/Cas9 components in the AeaNosC109^GD^ and AeaZpgC109^GD^ lines. GD lines (marked with mCherry) were balanced against a blocked GD (marked with eCFP) present at the CRISPR/Cas9 target site, and female trans-heterozygotes were then outcrossed to non-transgenic males. Average numbers of larvae +/- SEM and mean grouping are indicated for each group in the lower left-hand corner. Average and SEM superscripted with the same letter are not significantly different (p > 0.05). Total numbers of assessed larvae across all three replicates are indicated in the lower right corner. The embryos from this cross were subsequently injected with sgRNA targeting the TIMP-P4 locus, and the surviving larvae were reared to third instar, genotyped, and assayed for CRISPR/Cas9 activity at both the Carb109 locus (GD target) and the TIMP-P4 locus (exogenously applied sgRNA) via PCR and Sanger sequence trace analysis. The estimates of CRISPR/Cas9 activity were made using the Synthego ICE tool, which compares an edited trace to a non-transgenic trace.

### Testing for a parental contribution of CRISPR/Cas9 resulting in GD activity

To test for a parental effect resulting in GD activity, we performed reciprocal trans-heterozygous crosses as outlined above. In addition to female trans-heterozygotes outcrossed to non-transgenic males, we also tested male trans-heterozygotes outcrossed to non-transgenic females. In the first outcross (OX-1) of the trans-heterozygotes individuals, we found that the inheritance rates of the eCFP eye marker and the mCherry-marked GD were nearly 50% for both females and males (Figure 5) and were not statistically different when tested using pairwise T-testing (*p* = 0.5911 and *p* = 0.7587, respectively). When females from the eCFP-inheriting individuals were further outcrossed to non-transgenic (OX-2) to investigate the level of parental GD contribution, the median inheritance rate was 47% for offspring from eCFP expressing females and 49% for offspring from eCFP expressing male parentals, providing no evidence for a parental effect resulting in GD activity; regardless, a few data points were greater than 50% with two points reaching ∼80% (Figure 5).

**Figure 5.**
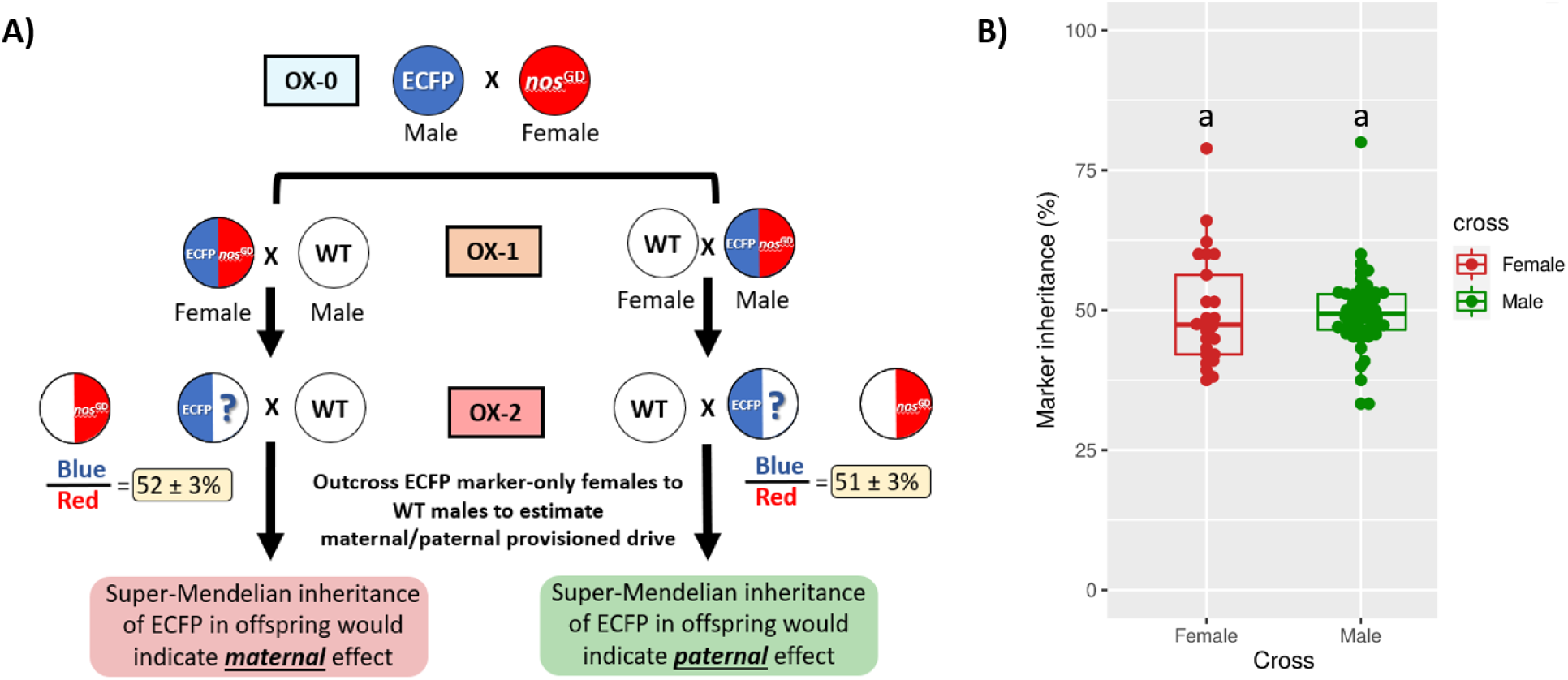
Locus balancing of an eCFP marker harboring line against the mCherry-marked AeaNosC109^GD^ line to test for parental GD effects. A) Crossing schematic for establishing trans-heterozygotes parentals (OX-1), which were then outcrossed to non-transgenic HWE mosquitoes to obtain the eCFP parentals for OX-2. The eCFP parentals used for OX-2 contained the eCFP marker in the Carb109 locus, allowing for drive into a non-transgenic allele if trans-acting Cas9-RNP was provisioned either maternally or paternally. B) eCFP marker inheritance for the final outcrossing of OX-2 to non-transgenic, where inheritance > 50% is suggestive of parentally supplied Cas9-RNP. Each data point in (B) represents the percentage of transgene inheritance resulting from the offspring of the second outcrossing of eCFP positive females to non-transgenic HWE males. The bar inside the box plot represents the median value, while the lower and upper borders of the boxes represent Q1 and Q3, respectively. Groups superseded by the same letter are not significantly different (p > 0.05). A minimum of 20 larvae was set for each group in order to be scored.

### Fitness of GD harboring mosquitoes and modeling of GD performance over multiple generations

Based on the experimentally derived data regarding GD inheritance including frequency of GDBI accrual, testing for maternal effect of the CRISPR/Cas9 RNP, and analysis of fitness parameters of the GD harboring mosquito lines (**Table S7**), we modeled the performance of AaeNosC109^GD^ or AeaZpgC109^GD^ in a hypothetical field release scenario for the purpose of population replacement (Figure 6; **Table S8**, **Table S9**). To model the lines, we used the MGDrivE mosquito population modeling package in R (41) and modeled for a static population of *Ae. aegypti* with a stable population size of 10,000 mosquitoes at a sex ratio of 1:1. We allowed for a single release of males at day 25 at a rate of 1/5^th^ of the population (*i.e.*, 2000 homozygous male mosquitoes) and ran 100 stochastic simulations for each of the transgenic lines until the allelic frequencies of the genotypes stabilized. Given that we targeted an intergenic region, all GBDI were assumed to incur no fitness cost to the mosquito, and we therefore merged the ‘B’ allele from MGDrivE (*i.e.,* a GD resistance allele that results in a fitness cost) with the ‘R’ allele (GD resistance allele with no fitness cost), such that all resistance alleles had the same fitness as the non-transgenic alleles (**Table S8**). Figure 6 displays the various prevalence levels of each genotype in the predicted model for females (all genotypes), males (all genotypes), and females divided between genotypes carrying at least one copy of the GD (displayed as “Hypothetical antiviral effector gene” to model a linked antiviral effector) and genotypes not carrying the GD (displayed as “Wildtype”). Overall, there was a rapid decline in the number of wildtype mosquitoes for both simulated releases and the rate at which the genotypes stabilized within a population differed between the two transgenic lines. In the AeaNosC109^GD^ line, the homozygous wildtype genotype was eliminated by ∼400 days-post release (PR) in the simulated closed system, with the hemizygous GD (no GDBI) and homozygous GD genotypes quickly replacing the homozygous wildtype genotype. By 450-500 days PR, however, the GDBI harboring resistance allele genotype became dominant and rapidly overtook the population eventually causing the GD to fall out of the population and the near entirety of the population to contain GDBI alleles by 1500 days PR. When a hypothetical antiviral effector linked to the GD was considered, the proportion of female mosquitoes carrying at least one copy of the antiviral effector gene peaked at 320 to 370 days PR with ∼80% of females carrying at least one copy of the antiviral effector. Any window of antiviral protection soon closed, and the GD was eliminated from the population by 1500 days PR. By comparison, the AeaZpgC109^GD^ line also eliminated the homozygous wildtype genotype by ∼400 days PR while GDBI resistance alleles became the dominant genotype in the population by ∼1000 days PR. Although the AeaZpgC109^GD^ was found to have lower GD inheritance rates than AeaNosC109^GD^, it was able to drive at least one copy of the GD through the population, reaching > 95% among all female mosquitoes by ∼420 days PR. Similar to the AeaNosC109^GD^ line, the window of protection for the hypothetically linked antiviral effector eventually closed due to the fitness differences between the GD and the GD-resistant (GDBI) alleles (**Table S9**), with > 50% of females losing the antiviral effector by ∼1500 days PR.

**Figure 6.**
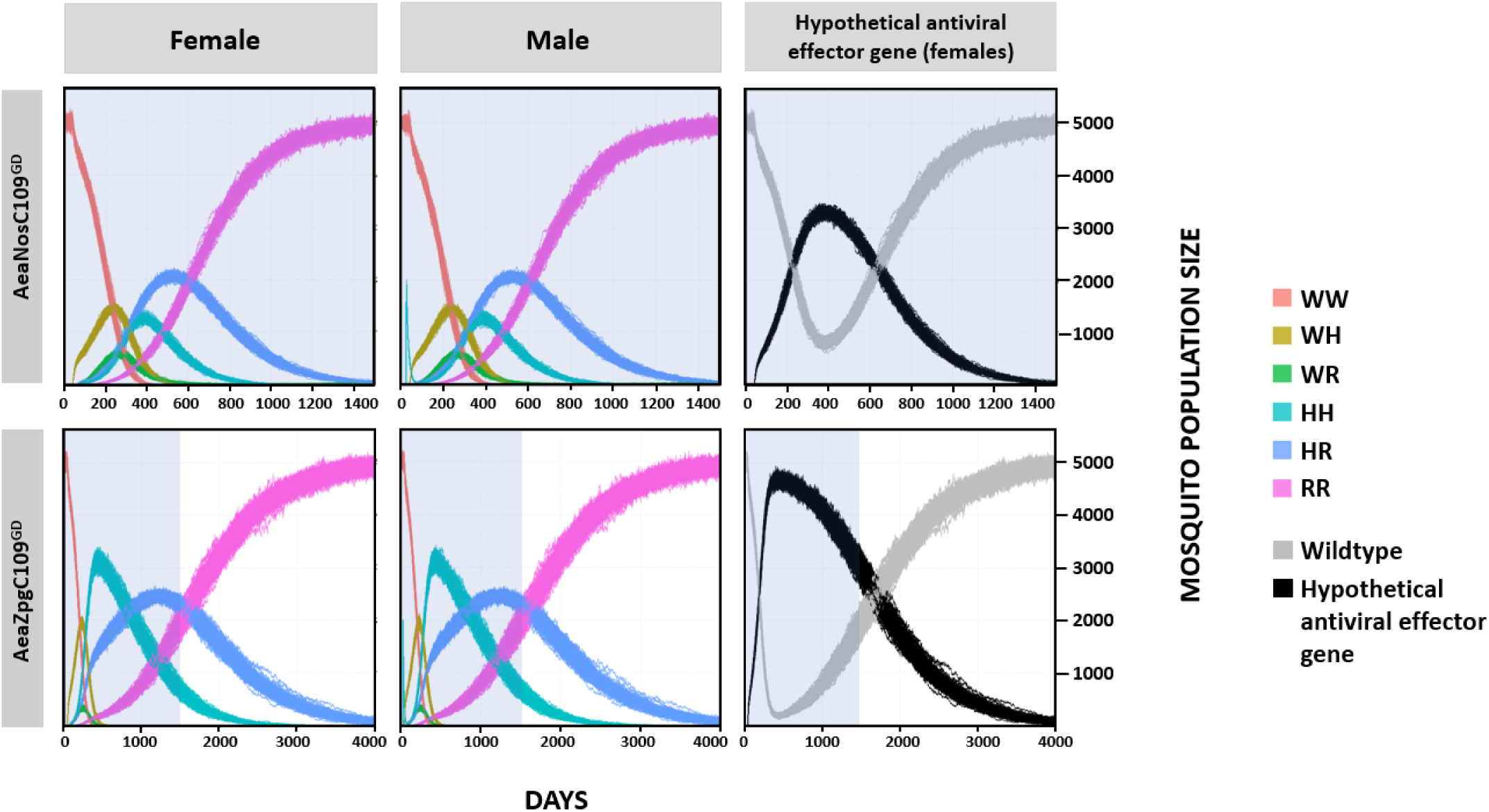
Population modeling for the AeaNosC109^GD^ line (upper panels) and the AeaZpgC109^GD^ line (lower panels) using MGDrivE. W = wildtype unedited allele, H = homing GD allele, R = GDBI resistance allele. A hypothetical refractory antiviral allele linked to the GD, shown as black in the right panels, displays the numbers of females in the population carrying at least one copy of the GD and a linked hypothetical ‘antiviral effector gene’. The time scale in days differs for the two transgenic lines with the pale blue backshading representing 1500 days post-release (PR) for comparison.

## DISCUSSION

Previous GD studies in mosquitoes have targeted recessive marker genes that provide for a visual phenotype when both alleles are disrupted. Since we selected intergenic loci, TIMP-P4 and Carb109, any disruption of these loci would not result in recessive phenotypes. We therefore positioned a codominant marker at the Carb109 locus, which then allowed us to distinguish between the GD cassette (mCherry marked) and a blocked drive (marked by eCFP). The CRISPR/Cas9 RNP requires two components: the Cas9 protein, and the sgRNA transcript. The Cas9 endonuclease was placed under control of a maternally-provisioned gene promoter, *nanos*, in the AeaNosC109^GD^ line or the germline-essential promoter for *innexin4* (*zpg*) in the AeaZpgC109^GD^ line (47, 52–53). Earlier work by Akbari *et al* (2013) indicated that both *nanos* and *innexin4* transcripts were detectable in the early embryo for up to 48 h, suggesting that *innexin4* (*zpg*) could be maternally-provisioned, a finding that was further supported based on observations from *zpg* knock-out studies in *Drosophila* (48, 54). In addition to Cas9, the active CRISPR/Cas9 RNP requires a sgRNA to guide the Cas9 protein for DNA cleavage. In our study, the expression of the sgRNA was under control of a snoRNA Pol-III promoter, which allowed for multiple possibilities of maternal inheritance of the CRISPR/Cas9 system. Assuming that Cas9 was maternally provisioned as either protein or mRNA, the possibilities of maternal inheritance included: 1) inheritance of the full Cas9-RNP complex, 2) inheritance of the Cas9 mRNA and sgRNA transcripts, 3) inheritance of Cas9 mRNA and/or protein, but not the sgRNA transcript. To test for this, we independently crossed the AeaNosC109^GD^ and AeaZpgC109^GD^ lines to the AeaeCFPC109 line to create females which, when mated to wildtype males, would have progeny that either inherited the GD or the null (non-editable) allele at the Carb109 locus.

The spatio-temporal expression of transgenes in *Ae. aegypti* is strongly influenced by the promoter used for expression as well as the genomic transgene insertion site, which can be affected by position effect variegation (55, 56). The importance of genomic position for optimal Cas9 activity in *Ae. aegypti* was identified in the work by Li *et al*. (2017), where the activity of Cas9 expression varied by genomic locus when placed under control of six different promoters and integrated into the *Ae. aegypti* genome in a quasi-random fashion using the *piggyBac* transposable element (29). The forward approach from Li *et al*. (2017) allowed for the identification of optimal genomic insertion loci for Cas9 activity in the germline, while in our study, we took a reverse approach by utilizing two genomic loci (TIMP-P4 and Carb109) previously identified to be highly permissive to antiviral transgene expression (13, 17). Our reasoning for this was that if the GD cassettes were positioned in a genomic locus that supported a strong expression of antiviral effectors, the entire system could be built and established as a single-component autonomous GD line. It was unclear, how an autonomous GD in an intergenic locus would perform. Our reverse engineering approach demonstrated that genomic loci that are well-suited towards the expression of antiviral effectors may not necessarily be ideal for the functioning of a CRISPR/Cas9 autonomous GD system. Notably, the presence of the GD system at the TIMP-P4 locus resulted in no GD activity and further demonstrated a very low level of inheritable GDBI when assessed via whole-amplicon sequencing. This suggests that the GD components were not efficiently expressed in the germline when positioned at the TIMP-P4 locus although our previous work demonstrated strong expression of transgenes in the adult female midgut at the same locus (17, 31). Meanwhile, a second genomic locus that had also been shown to be ideal for stable antiviral activity (*i.e.*, Carb109; [13]) exhibited high GD activity; however, there was a concomitant level of GDBI detectable that eventually blocked the GD in a population modeling simulation. This demonstrated that while the Carb109 locus supported GD activity by allowing rapid introgression of either GD cassette into the target population, the locus was prone to the generation of heritable GDBI, which over time blocked and eliminated the GD in the population modeling. These results suggest that when an autonomous CRISPR/Cas9 based GD is positioned at an intergenic locus, it will likely be self-exhausting. Our simulation models provided valuable information regarding the dynamics of the GD. For example, the AeaZpgC109^GD^ line provided a wider hypothetical window of protection (females harboring at least one copy of the GD and a hypothetical antiviral effector) than the AeaNosC109^GD^ line, although the latter GD line had stronger GD inheritance. The major differences in GD performance and biological parameters between the AeaNosC109^GD^ and AeaZpgC109^GD^ lines were the levels of GD activity, female deposition rate, resistance allele formation rate, and pupation success. To estimate which of these effects most strongly influenced the performance of the GD, we conducted simulations of the AeaNosC109^GD^ line in which maternal deposition rate, resistance allele formation rates, or pupation success parameters were substituted with the respective (experimentally determined) parameters from the AeaZpgC109^GD^ line (**Figure S4**). However, there was no appreciable difference in overall GD persistence when the levels of GDBI resistance alleles, or the levels of maternal deposition were reduced in the AeaNosC109^GD-SIMULATED^ line to those of the experimental AeaZpgC109^GD^ line (**Figure S4**). When the parameter of pupation success was modified to match that of the AeaZpgC109^GD^ line, the GD and associated hypothetical antiviral effector persisted through the population with approximately half of the females carrying at least one copy of the transgene up to ∼3000 days PR. These simulations suggest that while the rates of GD inheritance, GDBI resistance allele formation, and maternal deposition were important factors, it was the fitness of the GD line that had the greatest impact on the dynamics of GD inheritance. A study by Terradas *et al* (2021) investigated the deposition of germline expressed genes in the developing oocytes of autonomous *Anopheles* sp. GD lines (49), including the AgNosCd1^GD^ line (23, 49), which used the *nanos* promoter and 3’UTR for Cas9 expression. In their study, the authors found that *nanos* driven Cas9 transcripts were indeed expressed in the nurse cells along with native *nanos* transcripts; however, unlike the native *nanos* transcripts, the Cas9 transcripts were found at very low levels or were even absent in the developing oocyte (49). In males, by contrast, Cas9 expression controlled by the *nanos* regulatory elements paralleled that of the native *nanos* transcript. In regard to the *β2-tubulin* regulatory elements, it was shown by the same authors that in *An. stephensi, β2-tubulin* transcripts were male-specific. In addition, transcript expression occurred in the late stages of spermatogenesis, which may not be a suitable time window of expression when attempting to use the *β2-tubulin* promoter for a CRISPR/Cas9 based GD. Indeed, this could be a reason for the lack of GD activity that we observed for our Aeaβ2tC109^GD^ line. Furthermore, Terradas *et al* (2021) investigated the pattern of native *zpg* expression and found that its expression pattern was similar to that of *vasa* but at a lower general level. They found that there was expression of *zpg* throughout premeiotic and meiotic stages in males and high-level *zpg* expression in the sperm flagella, which could increase the prevalence of GDBI. In our AeaZpgC109^GD^ line, however, we identified a lower level of GDBI for male parentals and a lower level of GD compared to the AeaNosC109^GD^ line. Thus, it is likely that the expression of the Cas9 transcript under control of the *zpg* promoter does not perfectly parallel that of the native *zpg* transcript. Furthermore, although the *An. gambiae* AgNosCd1^GD^ line did not have an appreciable deposition of mRNA transcripts for Cas9 when expressed from the *nanos* promoter and 3’UTR (49), a recent analysis of GDBI by Carballar-Lejarazú *et al* (2022) found that there was indeed a maternal effect of CRISPR/Cas9 activity in the AgNosCd1^GD^ line (50). Taken together, these results suggest that the maternal contribution could be provisioned to the developing oocyte as residual Cas9 RNP complex might be expressed during early germline development and further persist in the zygote.

Simulations of the different GD performances and life parameters of the AeaNosC109^GD^ and AeaZpgC109^GD^ lines highlight the importance of fitness and maternal contribution with respect to the accrual of GDBI. Our population modeling revealed that in the absence of any fitness costs directly associated with GDBI, yet there was still a fitness cost associated with the GD as such because an autonomous GD in an intergenic locus will ultimately block itself due to the accrual of GDBI alleles and the drive will be lost. Therefore, an accumulation of GDBI alleles will ultimately exhaust and impair the GD. Given the significance of these observations obtained from our simulated release study, multi-generational cage trials aiming at modeling population replacement under more realistic field conditions (22, 23) will be imperative in the future to confirm the effects of fitness cost on single-component GD performance in *Ae. aegypti*.

Since we selected intergenic loci, which were assumed to have no effect on fitness if cleaved, the persistence of the autonomous drive system could be strengthened by its placement into a non-neutral (*e.g.*, coding region) locus as performed for *kh* in *An. stephensi* (21, 22). This could also extend the window of protection provided by the antiviral effector. There are environmental concerns regarding the release of transgenes into wild mosquito populations, and furthermore, there is a practical rationale for a non-persistent GD: the associated antipathogen effector gene could lose its efficacy over time. Such a loss of efficacy could be due to adaptations of the pathogen to evade the effect of the antipathogen effector gene, or spontaneous mutations leading to a loss of function of the antipathogen effector. An autonomous GD system that will ultimately self-block may have several benefits as it could rapidly drive an antipathogen transgene through wild mosquito populations while providing a window of protection from a circulating arbovirus, before dropping out of the population due to the fixation of GDBI alleles.

## DATA AVAILABILITY

The nucleotide sequences for all our GD constructs listed in Table S6 are available at NCBI (ncbi.nlm.nih.gov) under the following accession numbers. Construct AeaeCFPT4: accession # MT926371; construct AeaeCFPC109: accession # OL452018; construct AeaNosT4GD: accession # OL452014; construct Aeaβ2tC109GD: accession # OL452017; construct AeaNosC109GD: accession # OL452015; construct AeaZpgC109GD: accession # OL452016; construct pAeU6-MT: accession # OL452019; pAeT7ku70dsRNA: accession #: OL452021. The code modification to MGDrivE is provided in Table S8. Supplementary material is available at G3 online.

## ACKNOWLEDGEMENTS

We would like to thank Drs. Héctor Sánchez Castellanos, Rodrigo Corder, and John Marshall for their guidance and assistance with the use of the MGDrivE simulation for population modeling. We thank Drs. Hanno Schmidt and Gregory Lanzaro for kindly sharing the genomic vcf data for the *Ae. aegypti* genomes. Thanks to Susi Bennett for all her mosquito maintenance. We also thank the University of Missouri Genomics Technology Core for technical support.

## AUTHOR CONTRIBUTIONS

W.R., A.E.W., K.E.O., and A.W.E.F. designed research; W.R., A.E.W., J.L., I.S-V., and R.J. performed research; W.R. contributed new reagents/analytic tools; W.R., A.E.W., K.E.O., and A.W.E.F. analyzed data; and W.R., A.E.W., K.E.O., and A.W.E.F. wrote the paper.

