## Supplemental Data for "Assessing single-component gene drive systems in the mosquito *Aedes aegypti* via single generation crosses and modeling"

**This file includes:**

**Figure S1.** Size distributions of insertions/deletions for the NHEJ events observed in the three GD lines tested.
**Figure S2.** Schematic for the GD constructs used in our study**.**
**Figure S3.** Crossing schematic to test for maternal contributions of CRISPR/Cas9 components in the AeaNosC109$^{GD}$ and AeaZpgC109$^{GD}$ lines.
**Figure S4.** Modeling simulations for the AeaNosC109$^{GD}$ line (unmodified upper row) allowing for changed parameters for pupation success (xiF and xiM, second row), female deposition rate (dF, third row), and GDBI development rates (crF and crM, third row).

**Table S1.** Polymorphisms in the active target sites for the Carb109 and TIMP-P4 loci among 132 genomes of *Aedes aegypti*.
**Table S2.** Numbers of larval pools assessed and average metrics for gene drive inheritance for the OX-1 crosses.
**Table S3.** Numbers of larval pools assessed and average metrics for gene drive inheritance for the OX-2 crosses.
**Table S4.** Percentage of amplicons containing gene drive blocking indels (GDBI) for pooled negative larvae from the OX-2 generations.
**Table S5.** List of primers and gBlocks used in our study.
**Table S6.** NCBI sequences for the constructs used in our study.
**Table S7.** Life parameter data for hemizygote AeaNosC109$^{GD}$ and AeaZpgC109$^{GD}$ to assess fitness costs.
**Table S8.** Script modifications to the MGDrivE v1.6.0 Cube-CRISPR2MF.R to condense the B and R resistance alleles to a common no-fitness cost R allele. Original section of script taken from Sánchez et al (2020) (11).
**Table S9.** Fitness, GD, maternal deposition, and GDBI formation parameters used in the MGDrivE modeling for the AeaNosC109$^{GD}$ and AeaZpgC109$^{GD}$ lines.

**SI References**

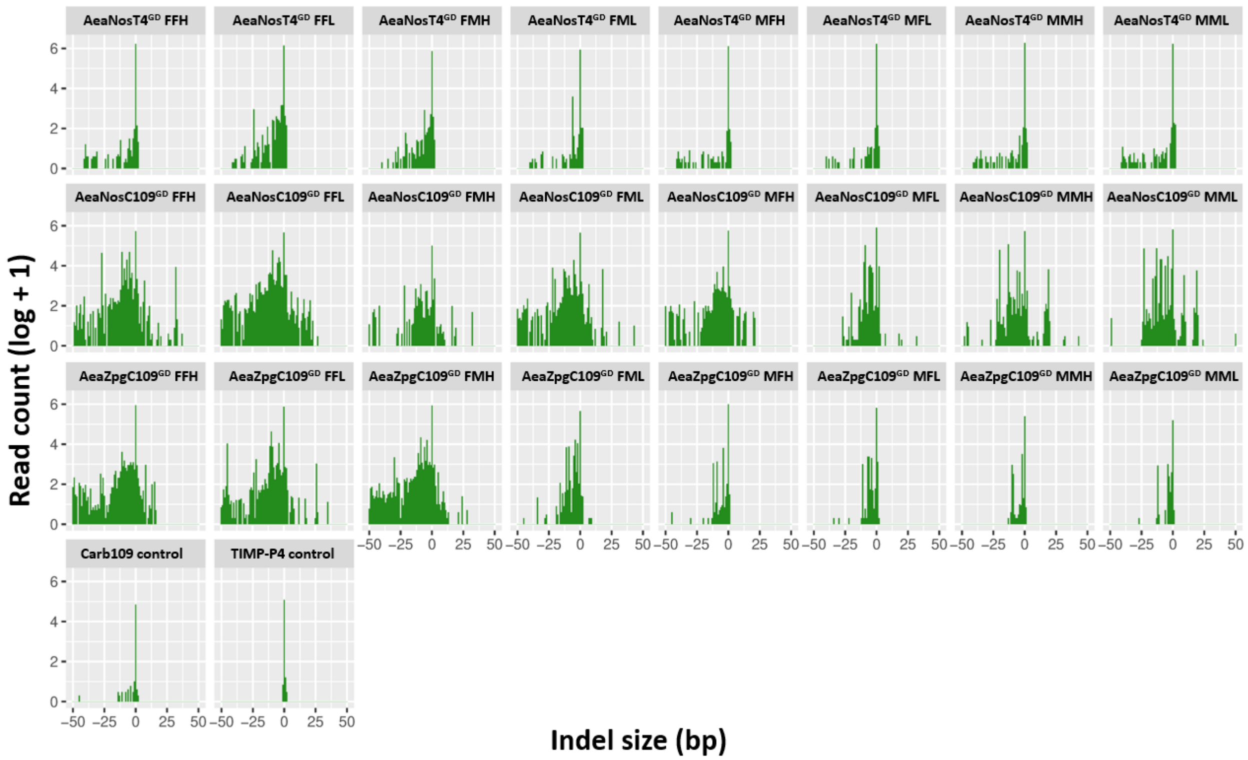


**Figure S1**. **Size distributions of insertions/deletions for the NHEJ events observed in the three GD lines tested.** The first letter within each cross indicates the parental transgenic sex, the second letter indicates the grandparental transgenic sex. L = low drive level in the parental generation, H = high drive in the parental generation. Control = Higgs' White Eye (HWE) non-transgenic mosquitoes. Data are presented as a log transformation.

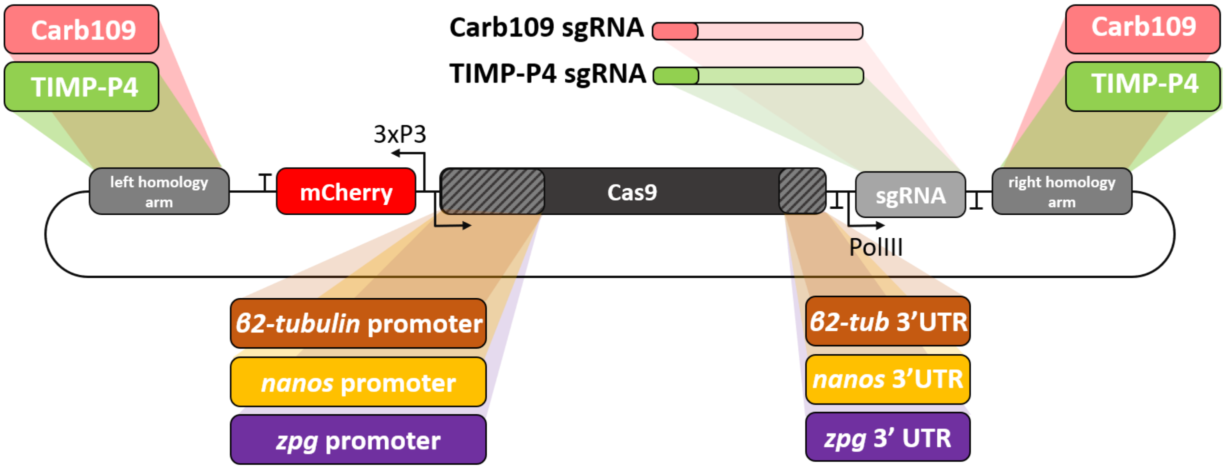


**Figure S2. Schematic for the GD constructs used in our study.** Word bubbles outside of the plasmid oval represent cassettes that were replaced within the construct to generate the five established transgenic lines tested in our study. Word bubbles that share matching coloration represent the cassettes used for each line; for example, for TIMP-P4, all of the upper green cassettes were contained within the same construct for insertion into the TIMP-P4 locus, while below, the yellow *nanos* promoter and nanos 3'-UTRs were used to drive Cas9 expression in the AeaNosT4$^{GD}$ line.

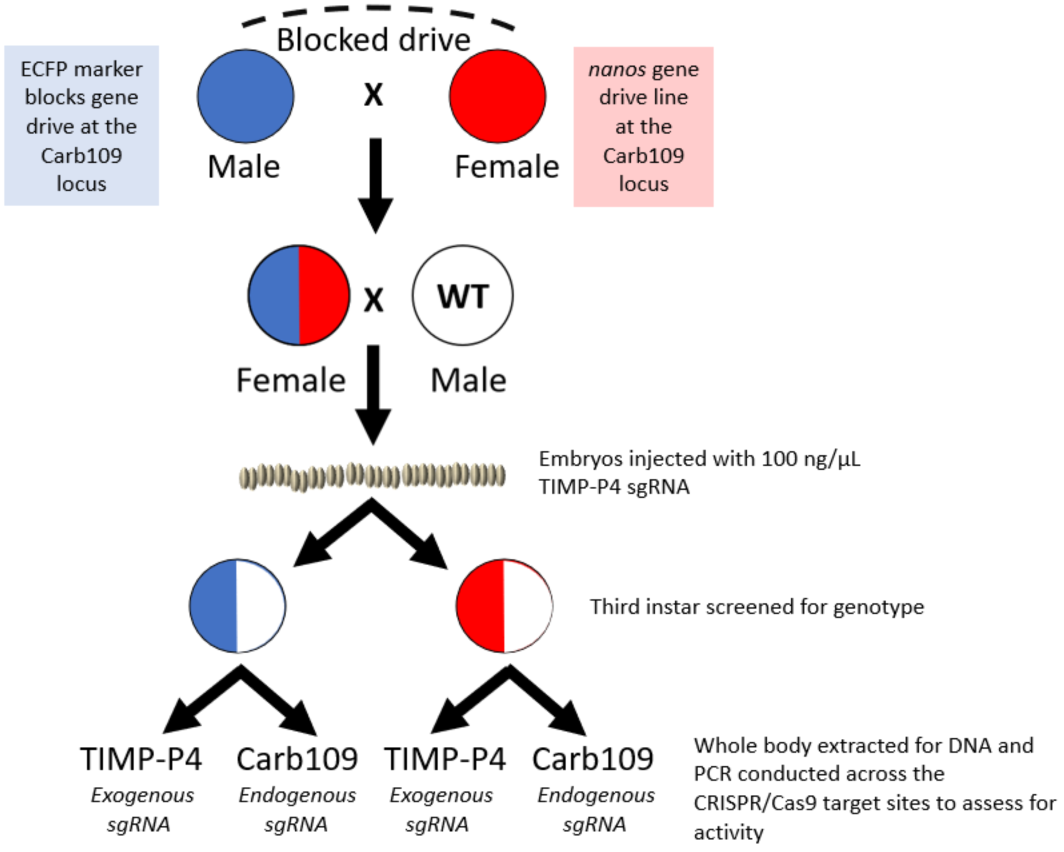


**Figure S3. Crossing schematic to test for maternal contributions of CRISPR/Cas9 components in the AeaNosC109$^{GD}$ and AeaZpgC109$^{GD}$ lines.** GD lines (mCherry marked) were balanced against a blocked GD (eCFP present at the CRISPR/Cas9 target site), and female trans-heeterozygotes were then outcrossed to non-transgenic males. The embryos from this cross were subsequently injected with sgRNA targeting the TIMP-P4 locus, and the surviving larvae were reared to third instar, genotyped, and assayed for CRISPR/Cas9 activity at both the Carb109 locus (GD target) and the TIMP-P4 locus (exogenously applied sgRNA) using PCR and Sanger sequence trace analysis.

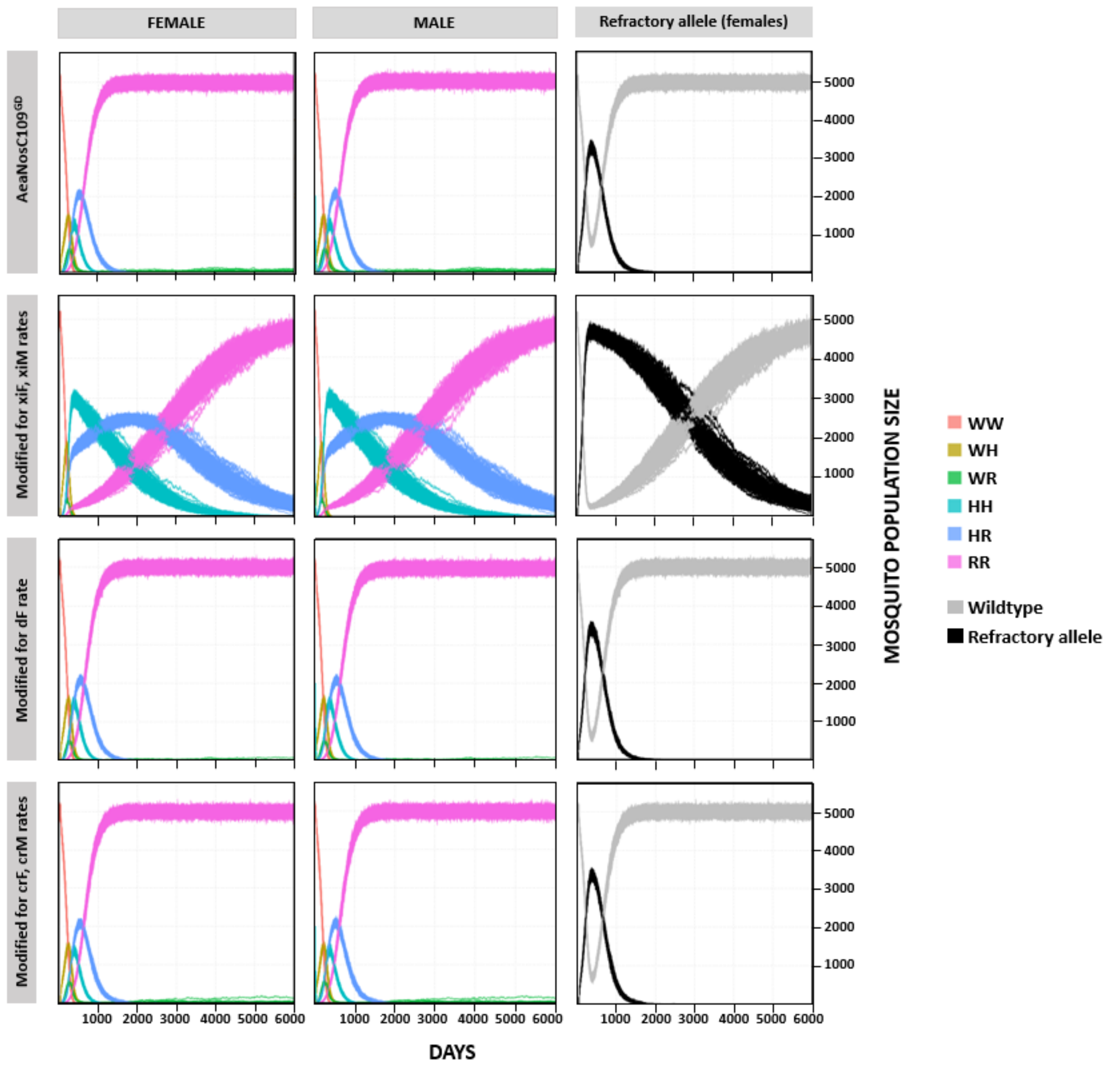


**Figure S4. Modeling simulations for the AeaNosC109$^{GD}$ line (unmodified upper row) using MGDrivE allowing for changed parameters for pupatory success (xiF and xiM, second row), female deposition rate (dF, third row), and GDBI development rates (crF and crM, third row).**

**Table S1. Polymorphisms in the active target sites for the Carb109 and TIMP-P4 loci among 132 genomes of *Aedes aegypti.*** The polymorphisms are indicated in boldface, while the PAM sequences are underlined. Data taken from Schmidt *et al*., 2020 (1).

| sgRNA | Polymorphisms (5' -protospacer + PAM -3') | Genomic locus* | Genome coverage | Polymorphism frequency |
|---|---|---|---|---|
| TIMP-P4 | GACCAACGGCAGTCATTGTG T**T**G | 2:321382218 | 230 | 0.026087 |
| | GACCAAC**C**GCAGTCATTGTG TGG | 2:321382231 | 222 | 0.00900901 |
| Carb109 | G**W**TATGCCGAAGAAAAGCCA GGG | 3:409699154 | 264 | 0.0113636 |

**Aedes aegypti* genome, strain Liverpool, version Vectorbase-54: LVP_AGWG.

**Table S2. Numbers of larval pools assessed and average metrics for GD inheritance among the OX-1 crosses.**

| Line | Cross | N groups | N total | Weighted average±SEM | Min | Max | N min | N max | Sample min | Sample max | Sample average (N) ±SEM |
|---|---|---|---|---|---|---|---|---|---|---|---|
| AeaNosT4$^{GD}$ | Female | 22 | 1131 | 46.6±5.3 | 40.3 | 56.3 | 67 | 64 | 22 | 106 | 51.4±4.8 |
| | Male | 24 | 2072 | 48.7±5.2 | 27.9 | 59.7 | 43 | 57 | 22 | 182 | 86.3±8.7 |
| AeaNosC109$^{GD}$ | Female | 20 | 1380 | 73.3±12.3 | 48.7 | 94.0 | 37 | 67 | 23 | 142 | 69.0±6.1 |
| | Male | 20 | 1864 | 70.1±15.8 | 45.7 | 96.1 | 81 | 77 | 23 | 172 | 93.2±7.3 |
| Aeaβ2tC109$^{GD}$ | Female | 10 | 409 | 48.2±4.0 | 43.4 | 55.0 | 53 | 20 | 20 | 63 | 40.9±4.6 |
| | Male | 15 | 1029 | 47.5±2.7 | 42.9 | 55.2 | 56 | 29 | 29 | 142 | 68.6±8.8 |
| AeaZpgC109$^{GD}$ | Female | 13 | 711 | 66.1±9.8 | 48.3 | 80.4 | 58 | 51 | 34 | 77 | 54.7±4.0 |
| | Male | 19 | 1807 | 55.6±8.3 | 42.3 | 77.8 | 97 | 27 | 21 | 174 | 95.1±8.9 |

**Table S3. Numbers of larval pools assessed and average metrics for GD inheritance among the OX-2 crosses.**

| Line | Cross | N groups | N total | Weighted average±SEM | Min | Max | N min | N max | Sample min | Sample max | Sample (N) average±SEM |
|---|---|---|---|---|---|---|---|---|---|---|---|
| AeaNosT4$^{GD}$ | FFH | 26 | 1451 | 49.4±18.9 | 33.3 | 89.5 | 21 | 124 | 21 | 124 | 55.8±4.9 |
| | FFL | 29 | 1441 | 47.7±6.4 | 38.1 | 60.0 | 21 | 25 | 21 | 113 | 49.7±4.6 |
| | FMH | 23 | 795 | 48.4±8.9 | 40.0 | 72.0 | 25 | 25 | 21 | 58 | 34.6±2.5 |
| | FML | 24 | 1219 | 47.4±7.4 | 34.8 | 54.2 | 23 | 59 | 23 | 110 | 50.8±3.8 |
| | MFH | 31 | 1894 | 46.5±9.0 | 26.6 | 55.3 | 79 | 47 | 25 | 98 | 61.1±4.1 |
| | MFL | 30 | 2350 | 48.1±6.6 | 39.2 | 68.4 | 51 | 57 | 24 | 156 | 78.3±5.2 |
| | MMH | 27 | 1735 | 48.8±5.3 | 38.6 | 60.0 | 44 | 20 | 20 | 120 | 64.3±5.8 |
| | MML | 28 | 2521 | 49.2±4.7 | 37.9 | 58.6 | 66 | 29 | 22 | 195 | 90.0±10.3 |
| AeaNosC109$^{GD}$ | FFH | 28 | 1971 | 68.4±29.0 | 43.0 | 91.4 | 107 | 140 | 20 | 151 | 70.4±7.3 |
| | FFL | 29 | 2032 | 66.6±32.4 | 43.5 | 100.0 | 69 | 114 | 29 | 114 | 70.1±4.7 |
| | FMH | 22 | 1775 | 69.7±28.1 | 46.1 | 95.9 | 89 | 49 | 23 | 200 | 80.7±8.0 |
| | FML | 27 | 1690 | 57.4±24.0 | 22.9 | 93.9 | 35 | 65 | 22 | 135 | 62.6±6.9 |
| | MFH | 27 | 2287 | 65.9±29.8 | 40.0 | 100.0 | 105 | 44 | 26 | 223 | 84.7±11.2 |
| | MFL | 26 | 2636 | 76.4±37.0 | 42.9 | 100.0 | 77 | 137 | 24 | 237 | 101.4±12.3 |
| | MMH | 28 | 2734 | 72.7±35.4 | 36.8 | 100.0 | 76 | 20 | 20 | 232 | 97.6±9.8 |
| | MML | 32 | 3162 | 62.4±25.7 | 36.0 | 98.1 | 75 | 53 | 20 | 212 | 98.8±9.6 |
| AeaZpgC109$^{GD}$ | FFH | 16 | 1129 | 56.9±9.9 | 40.0 | 78.8 | 40 | 33 | 20 | 167 | 70.6±10.4 |
| | FFL | 21 | 1413 | 58.5±21.6 | 30.4 | 89.2 | 23 | 37 | 22 | 164 | 67.3±10.3 |
| | FMH | 22 | 1651 | 58.7±13.5 | 42.1 | 79.3 | 38 | 87 | 27 | 176 | 75.1±8.5 |
| | FML | 16 | 1017 | 58.4±17.2 | 34.8 | 79.7 | 23 | 59 | 23 | 137 | 63.6±8.2 |
| | MFH | 20 | 1348 | 56.7±19.4 | 39.5 | 84.7 | 86 | 111 | 21 | 174 | 67.4±9.2 |
| | MFL | 20 | 1531 | 59.4±19.5 | 39.8 | 90.9 | 103 | 22 | 21 | 179 | 76.6±10.4 |
| | MMH | 23 | 1925 | 54.2±21.7 | 27.7 | 91.9 | 137 | 74 | 20 | 176 | 83.7±11.3 |
| | MML | 19 | 1308 | 52.2±25.7 | 19.1 | 81.9 | 21 | 177 | 21 | 177 | 68.8±10.9 |

**Table S4. Percentage of amplicons containing gene drive blocking indels (GDBI) for pooled negative larvae from the OX-2 generations.**

| Line | Cross | N groups | Median (%) | Q1 (%) | Q3 (%) | Min (%) | Max (%) |
|---|---|---|---|---|---|---|---|
| AeaNosT4$^{GD}$ | FFH | 10 | 0.020 | 0.017 | 0.025 | 0.01 | 0.07 |
| AeaNosT4$^{GD}$ | FFL | 11 | 0.021 | 0.019 | 0.026 | 0.01 | 4.2 |
| AeaNosT4$^{GD}$ | FMH | 11 | 0.117 | 0.051 | 0.54 | 0.02 | 1.5 |
| AeaNosT4$^{GD}$ | FML | 10 | 0.040 | 0.025 | 0.158 | 0.01 | 2.8 |
| AeaNosT4$^{GD}$ | MFH | 10 | 0.018 | 0.016 | 0.022 | 0.01 | 0.03 |
| AeaNosT4$^{GD}$ | MFL | 10 | 0.021 | 0.019 | 0.024 | 0.01 | 0.03 |
| AeaNosT4$^{GD}$ | MMH | 11 | 0.019 | 0.015 | 0.024 | 0.01 | 0.03 |
| AeaNosT4$^{GD}$ | MML | 12 | 0.022 | 0.017 | 0.026 | 0.01 | 0.55 |
| AeaNosC109$^{GD}$ | FFH | 10 | 19.1 | 4.9 | 32.33 | 0.71 | 98.1 |
| AeaNosC109$^{GD}$ | FFL | 9 | 12.4 | 11.5 | 22.24 | 6.0 | 53.9 |
| AeaNosC109$^{GD}$ | FMH | 5 | 4.7 | 2.4 | 10.26 | 0.79 | 25.9 |
| AeaNosC109$^{GD}$ | FML | 9 | 11.9 | 8.0 | 18.15 | 0.06 | 26.5 |
| AeaNosC109$^{GD}$ | MFH | 9 | 0.15 | 0.06 | 1.70 | 0.02 | 14.8 |
| AeaNosC109$^{GD}$ | MFL | 10 | 9.2 | 2.8 | 25.23 | 0.19 | 68.0 |
| AeaNosC109$^{GD}$ | MMH | 9 | 1.6 | 0.16 | 63.94 | 0.03 | 92.0 |
| AeaNosC109$^{GD}$ | MML | 10 | 14.1 | 0.36 | 38.39 | 0.02 | 100.0 |
| AeaZpgC109$^{GD}$ | FFH | 8 | 2.1 | 1.1 | 3.48 | 0.02 | 4.5 |
| AeaZpgC109$^{GD}$ | FFL | 11 | 2.6 | 1.5 | 17.87 | 0.09 | 53.9 |
| AeaZpgC109$^{GD}$ | FMH | 11 | 2.0 | 0.56 | 12.15 | 0.03 | 21.5 |
| AeaZpgC109$^{GD}$ | FML | 5 | 11.9 | 2.9 | 13.37 | 0.03 | 14.2 |
| AeaZpgC109$^{GD}$ | MFH | 9 | 0.04 | 0.03 | 0.19 | 0.02 | 3.8 |
| AeaZpgC109$^{GD}$ | MFL | 8 | 1.6 | 0.04 | 2.04 | 0.01 | 22.7 |
| AeaZpgC109$^{GD}$ | MMH | 11 | 0.02 | 0.01 | 1.46 | 0.01 | 6.7 |
| AeaZpgC109$^{GD}$ | MML | 9 | 0.07 | 0.01 | 0.63 | 0.00 | 14.6 |
| HWE | Carb109 | 1 | 0.04% | | | | |
| HWE | TIMP-P4 | 1 | 0.02% | | | | |

**Table S5. List of primers and gBlocks used in our study.**

| Primer | Name | Sequence (5’ -> 3’) |
| --- | --- | --- |
| BR-26 | AAEL017774PROMF | CCATCTAGAGACTGGTCTTTTAGTTATGACTTCTG |
| BR-27 | AAEL017774PROMR | TTCGGATCCGAAGACCCATTTCACTACTCTTGCCTCTGC |
| BR-28 | AAEL019894_3pUTRF-SalI | CATGTCGACAAGCCTAGAAGGAATAAAAATGGCTGAA ATTATCCG |
| BR-29 | AAEL019894_3pUTRR-XbaI | GCATCTAGAATCACTCTTACGACACTGCATAC |
| BR-32 | AAEL019894PF_HindIII | CATAAGCTTAAACTCGAGTCGTGTGGCAAGTACGAAG |
| BR-33 | AAEL019894PR_PstI-NcoI | CATCTGCAGCCATGGTGGAGCACTTCTAGCGGTTC |
| BR-34 | AG45945F-NcoI | CATCCATGGACTATAAGGACCACGACG |
| BR-34 | AG45945F-NcoI | CATCCATGGACTATAAGGACCACGACG |
| BR-35 | AG45945R-SalI | CATGTCGACTTACTTTTTCTTTTTTGCCTGGCC |
| BR-35 | AG45945R-SalI | CATGTCGACTTACTTTTTCTTTTTTGCCTGGCC |
| BR-40 | ZA_AeNosPR-XhoIF | CATCTCGAGCACTATCAAACCCCTAAAGACA |
| BR-41 | ZA_AeNosPR-PciIR | CATCTCGAGACATGTTGTTCGTTGATCTCGATCAGCCA |
| BR-42 | ZA_AeNos3pU-StuIF | CATAGGCCTCGTAATCGAAGTGTTGGACGGGGAAAG |
| BR-43 | ZA_AeNos3pU-NheIR | CATGCTAGCACCAGACATCCGTTTAAGCTGAACCAC |
| BR-44 | ku70-ZA2341T7-F | TAATACGACTCACTATAGGGTTTCGCTGGGTGTCAACA TTTATTCC |
| BR-45 | ku70-ZA2342T7-R | TAATACGACTCACTATAGGGCTTGATTTTCTTCGCTACT GACTTGA |
| BR-54 | 3xP3FOR_XhoIF | CATCTCGAGAAATTCGAGCTCGCCCGG |
| BR-55 | SV40REV_XhoIR | CATCTCGAGCCGTACGCGTATTCGATAAG |
| BR-60 | Carb109scrF_3161 | CGCACCTAATCAGACAGTCG |
| BR-80 | T7_dsRNA_HindIII | CATAAGCTTAATACGACTCACTATAGGG |
| BR-98 | SV40_GibREV | AGCGGCCGCGACTCTAGA |
| BR-99 | 3xP3_GibFOR | GGTGGCGACCGGTGGATC |
| BR-100 | TIMP4donBK-R | GATCTAGAGTCGCGGCCGCTTTACTTGTACAGCTCGTC CATG |
| BR-101 | TIMP4donBK-F | ACGATCCACCGGTCGCCACCATGGTGAGCAAGGGCGA G |
| BR-102 | Blac-GibFOR | CTGCTATGTGGCGCGGTATTATC |
| BR-103 | Blac_GibREV | GATAATACCGCGCCACATAGCAG |
| BR-112 | Carb109_g15LAHR | CAATCTCGAGCCAGGGTCAGGGAGTTGGA |
| BR-113 | Carb109_g15RAHF | CAATCTCGAGCTTTTCTTCGGCATATCTATCGCTAGG |
| BR-349 | U6empty_Gblock | TCCACATCTCCATACATTCAACGCACTGTGCGGCTGTGC TGTGCGACTCCGTCGAGTCGACCAACATAGTTGAAACA AATTGAATATTTAATTGATCGTTATAGGAATGGTGTTA GATGAGTCATCCTTTACAGTAAGCACATACAGTATTATA ATTGAAGATCGTCGGCAGATAGGTGTGTAGGGTAGAG TATCAGCAATAAGTTGGGACGTTTGACTTTTTGTAGGT AGACAAAAACTAAACTTTTTTTCGCTTCTCTATGTGTGC CCCCCCGGGTAGCGTATCGTTCCGATTGTGGTGCGAAC GAATGAAATCGCCTATCGAGTTGATACGTCCATCTATC GCTAGAACCGCGTTCGCTGTAAAAGACTATATAAGAGC AGAGGCAAGAGTAGTGAAATGGGTCTTCGAGAAGACC TGTTTCAGAGCTATGCTGGAAACAGCATAGCAAGTTGA |

| | | |
|---|---|---|
| | | AATAAGGCTAGTCCGTTATCAACTTGAAAAAGTGGCAC CGAGTCGGTGCTTTTTTTTTTTT |
| BR-350 | SacII-U6MT-F | CAATATCCGCGGTCCACATCTCCATACATTCAACG |
| BR-351 | SacII-U6MT-R | TAGATGCCGCGGAAAAAAAAAAAAAAAAGCACCGACTC GGTGCCAC |
| BR-360 | TIMP4-Pstop | AAATGGACCAACCGCAGTCATTGTG |
| BR-361 | TIMP4-Psbot | AAACCACAATGACTGCGGTTGGTCC |
| BR-362 | C109-Pstop | AAATGGATATGCCGAAGAAAAGCCA |
| BR-363 | C109-Psbot | AAACTGGCTTTTCTTCGGCATATCC |
| BR-364 | C109RAH-SacII-R | CAATACCGCGGATGGGATGCAGAACCATTTG |
| BR-368 | KpnI-Carb109scrF_3161 | CAATAGGTACCGCACCTAATCAGACAGTCG |
| BR-655 | zpgPROM-R_BsbI-PstI | CATTATCTGCAGGAAGACCCGATGATTTAGGGGTTTG |
| BR-656 | zpgF_131-XhoI | CAATACTCGAGATGAATCCTAAAGTCCTGCTCG |
| BR-660 | zpg3PUTR-F_BbsI-PstI | AAGTAACTGCAGGAAGACCTTCGATAAAAGTATCGTCC TAAGACTTATTCAG |
| BR-666 | Cas9F-GG-ZPGredo | CAATATGAAGACGGCATCATGGACTATAAGGACCACG ACG |
| BR-667 | Cas9R-GG-ZPGredo | CAATATGAAGACGGTCGATTACTTTTTCTTTTTTGCCTG GCCG |
| BR-683 | inx4-GIB_Rev | TGGACATGCATTAGACTTACTAGTAGGTGTTTGGACCA AGTGGAG |
| BR-724 | C109NGS-F | ACACTCTTTCCCTACACGACGCTCTTCCGATCTCGCACC TAATCAGACAGTCG |
| BR-725 | C109NGS-R | GTGACTGGAGTTCAGACGTGTGCTCTTCCGATCTCCTG CCTTCATTAAGCTCTTTG |
| BR-726 | TIMPNGS-F | ACACTCTTTCCCTACACGACGCTCTTCCGATCTAACGAG ATGCCTTCTCCTGA |
| BR-727 | TIMPNGS-R | GTGACTGGAGTTCAGACGTGTGCTCTTCCGATCTAAAA TGGCGTTCGATGAGA |

**Table S6**. **NCBI sequences for the constructs used in our study.**

| Construct | Purpose | NCBI accession |
|---|---|---|
| AeaeCFPT4 | Test knock-in plasmid for the TIMP-P4 locus | MT926371 |
| AeaeCFPC109 | Test knock-in plasmid for the Carb109 locus | OL452018 |
| AeaNosT4$^{GD}$ | Gene drive line for nanos Cas9 in the TIMP-P4 locus | OL452014 |
| Aeaβ2tC109$^{GD}$ | Gene drive line for β2tubulin Cas9 in the Carb109 locus | OL452017 |
| AeaNosC109$^{GD}$ | Gene drive line for nanos Cas9 in the Carb109 locus | OL452015 |
| AeaZpgC109$^{GD}$ | Gene drive line for zpg Cas9 in the Carb109 locus | OL452016 |
| pAeU6-MT | AAEL017774 promoter with inverted *Bbs*I sites and chiRNA (Dang *et al*., 2015) sgRNA (5) | OL452019 |
| pAeT7ku70dsRNA | Template plasmid for the anti-ku70 dsRNA trigger from Basu *et al*. (2015) (7) flanked by T7 RNA promoters. | OL452021 |

**Table S7. Life parameter data for hemizygote AeaNosC109$^{GD}$ and AeaZpgC109$^{GD}$ to assess fitness costs.** Data derived from hemizygous males or HWE males that were allowed to mate with female HWE. * = 0.01 < $p$ < 0.05 compared to HWE by one-tailed t-test. No stars indicate non-significant difference by one-way ANOVA.

| Line | WT (HWE) | AeaNosC109$^{GD}$ | AeaZpgC109$^{GD}$ |
|---|---|---|---|
| larva-to-pupa development (in days) | 9.8 ± 4.3 | 9.5 ± 5.6 | 9.5 ± 7.0 |
| larva viability (% survival) | 44.2 ± 7.3 | 30.9 + 2.9* | 44.8 ± 2.5 |
| % female adults | 45.4 ± 1.6 | 47.8 ± 1.5 | 48.2 ± 1.1 |
| % male adults | 54.6 ± 1.7 | 52.2 ± 1.5 | 51.8 ± 1.1 |
| 50% female survival (days) | 34 ± 6.2 | 44.5 | 41.5 |
| 50% male survival (days) | 20.7 ± 3.3 | 16 ± 1.6 | 18 ± 4.5 |
| fecundity (# eggs) | 62.6 ± 2.8 | 57.6 ± 2.6 | 64.6 ± 2.0 |
| fertility (egg hatchability) (%) | 49.2 ± 8.2 | 44.0 ± 3.6 | 58.3 ± 3.4 |
| male contribution (# pos / total n) | | 35.5 ± 1.2<br>(1931/5447) | 25.6<br>(1035/4049) |

**Table S8. Script modifications to the MGDrivE v1.6.0 Cube-CRISPR2MF.R to condense the B and R resistance alleles to a common no-fitness cost R allele.** Original section of script taken from Sánchez *et al.* (2020) (2).

```
##REMOVED B GENOTYPES IN THE GTYPE VECTOR
 ## define matrices
 ## Matrix Dimensions Key: [femaleGenotype,maleGenotype,offspringGenotype]
 gtype <- c('WW', 'WH', 'WR', 'HH', 'HR', 'RR')
 size <- length(gtype)
 tMatrix <- array(data=0, dim=c(size, size, size), dimnames=list(gtype, gtype, gtype))
#transition matrix

 ## fill tMatrix with probabilities
 ## COMMENTED OUT ANY CROSSES WITH A B ALLELE AND THEN REDID ANYTHING IN
THE PROBABILITIES THAT WOULD GIVE A B ALLELE
 #('WW', 'WH', 'WR', 'HH', 'HR', 'RR')
 tMatrix['WW','WW', 'WW'] <- 1

 tMatrix['WR','WW', c('WW', 'WR')] <- c( 1, 1)/2
 tMatrix['WR','WR', c('WW', 'WR', 'RR')] <- c( 1/2, 1, 1/2)/2

 tMatrix['HH','HH', 'HH'] <- 1

 tMatrix['HR','HH', c('HH', 'HR')] <- c( 1, 1)/2
 tMatrix['HR','HR', c('HH', 'HR', 'RR')] <- c( 1/2, 1, 1/2)/2

 tMatrix['RR','WW', 'WR'] <- 1
 tMatrix['RR','WR', c('WR', 'RR')] <- c( 1, 1)/2
 tMatrix['RR','HH', 'HR'] <- 1
 tMatrix['RR','HR', c('HR', 'RR')] <- c( 1, 1)/2
 tMatrix['RR','RR', 'RR'] <- 1

 ## set the other half of the matrix that is symmetric
 # Boolean matrix for subsetting, used several times
 boolMat <- upper.tri(x = tMatrix[ , ,1], diag = FALSE)
 # loop over depth, set upper triangle
 for(z in 1:size){tMatrix[ , ,z][boolMat] <- t(tMatrix[ , ,z])[boolMat]}


 ## fill asymmetric parts of tMatrix
 #female specific homing, except for WHxWH

 tMatrix['WH','WW',] <- c((1-cF)*(1-dF), (1+cF*chF)*(1-dF), ((1-cF)*dF*drF + (cF*(1-
chF)*crF)*(1-dF)) +((1-cF)*dF*(1-drF) + (cF*(1-chF)*(1-crF))*(1-dF)),
```

```
(1+cF*chF)*dF*dhF, ((1+cF*chF)*dF*(1-dhF)*drF) + ((1+cF*chF)*dF*(1-dhF)*(1-drF)),
((cF*(1-chF)*crF)*dF*drF)+((cF*(1-chF)*crF)*dF*(1-drF) + (cF*(1-chF)*(1-
crF))*dF*drF)+((cF*(1-chF)*(1-crF))*dF*(1-drF)))/2

 tMatrix['WH','WH',] <- c((1-cF)*(1-cM)*(1-dF),
                        (1+cF*chF)*(1-cM)*(1-dF) + (1-cF)*(1+cM*chM),
                        ((1-cF)*(1-cM)*dF*drF + cF*(1-chF)*crF*(1-cM)*(1-dF) + (1-cF)*cM*(1-
chM)*crM)+((1-cF)*(1-cM)*dF*(1-drF) + cF*(1-chF)*(1-crF)*(1-cM)*(1-dF) + (1-
cF)*cM*(1-chM)*(1-crM)),
                        (1+cF*chF)*(1-cM)*dF*dhF + (1+cF*chF)*(1+cM*chM),
                        ((1+cF*chF)*(1-cM)*dF*(1-dhF)*drF + cF*(1-chF)*crF*(1+cM*chM) +
(1+cF*chF)*cM*(1-chM)*crM)+((1+cF*chF)*(1-cM)*dF*(1-dhF)*(1-drF) + cF*(1-
chF)*(1-crF)*(1+cM*chM) + (1+cF*chF)*cM*(1-chM)*(1-crM)),
                        (cF*(1-chF)*crF*(1-cM)*dF*drF + cF*(1-chF)*crF*cM*(1-
chM)*crM)+(cF*(1-chF)*crF*(1-cM)*dF*(1-drF) + cF*(1-chF)*(1-crF)*(1-cM)*dF*drF +
cF*(1-chF)*(1-crF)*cM*(1-chM)*crM + cF*(1-chF)*crF*cM*(1-chM)*(1-crM))+(cF*(1-
chF)*(1-crF)*(1-cM)*dF*(1-drF) + cF*(1-chF)*(1-crF)*cM*(1-chM)*(1-crM)))/4

 tMatrix['WH','WR', ] <- c((1-cF)*(1-dF), (1+cF*chF)*(1-dF), ((1-cF)*dF*drF + (cF*(1-
chF)*crF)*(1-dF) + 1-cF) + ((1-cF)*dF*(1-drF) + (cF*(1-chF)*(1-crF))*(1-dF)),
(1+cF*chF)*dF*dhF, ((1+cF*chF)*dF*(1-dhF)*drF + 1+cF*chF ) + ((1+cF*chF)*dF*(1-
dhF)*(1-drF)), ((cF*(1-chF)*crF)*dF*drF + cF*(1-chF)*crF) + ((cF*(1-chF)*crF)*dF*(1-
drF) + (cF*(1-chF)*(1-crF))*dF*drF + cF*(1-chF)*(1-crF)) + ((cF*(1-chF)*(1-crF))*dF*(1-
drF)))/4

  tMatrix['WH','RR',c('WR', 'HR', 'RR')] <- c(1-cF, 1+cF*chF, (cF*(1-chF)*crF)+(cF*(1-
chF)*(1-crF)))/2

 tMatrix['WH','HH',c('WH', 'HH', 'HR')] <- c(1-cF, 1+cF*chF, (cF*(1-chF)*crF)+(cF*(1-
chF)*(1-crF)))/2

 tMatrix['WH','HR',c('WH', 'HH', 'HR',
 'WR', 'RR')] <- c(1-cF, 1+cF*chF, (cF*(1-chF)*crF + 1+cF*chF)+(cF*(1-chF)*(1-crF)),
 1-cF, (cF*(1-chF)*crF)+(cF*(1-chF)*(1-crF)))/4

 # female deposition things
 tMatrix['HH','WW', c('WH', 'HH', 'HR')] <- c( 1-dF, dF*dhF, (dF*(1-dhF)*drF)+(dF*(1-
dhF)*(1-drF)))

 tMatrix['HH','WR', c('WH', 'HH', 'HR')] <- c( 1-dF, dF*dhF, (dF*(1-dhF)*drF + 1)
+(dF*(1-dhF)*(1-drF)))/2

  tMatrix['HR','WW', c('WH', 'WR', 'HH', 'HR',
```

```
 'RR')] <- c( 1-dF, 1-dF, dF*dhF, (dF*(1-dhF)*drF)+(dF*(1-dhF)*(1-drF)),
(dF*drF)+(dF*(1-drF)))/2
 tMatrix['HR','WR', c('WH', 'WR', 'HH', 'HR',
 'RR')] <- c( 1-dF, 1-dF, dF*dhF, (dF*(1-dhF)*drF + 1)+(dF*(1-dhF)*(1-drF)), (dF*drF +
1)+(dF*(1-drF)))/4

 #male specific homing

 tMatrix['WW','WH', c('WW', 'WH', 'WR')] <- c(1-cM, 1+cM*chM, (cM*(1-
chM)*crM)+(cM*(1-chM)*(1-crM)))/2
 tMatrix['WR','WH', c('WW', 'WH', 'WR',
 'HR', 'RR')] <- c(1-cM, 1+cM*chM, (cM*(1-chM)*crM + 1-cM)+(cM*(1-chM)*(1-crM)),
 1+cM*chM, (cM*(1-chM)*crM)+(cM*(1-chM)*(1-crM)))/4
 tMatrix['HH','WH', c('WH', 'HH',
 'HR')] <- c((1-cM)*(1-dF), 1+cM*chM + (1-cM)*dF*dhF,
 (cM*(1-chM)*crM + (1-cM)*dF*(1-dhF)*drF)+(cM*(1-chM)*(1-crM) + (1-cM)*dF*(1-
dhF)*(1-drF)))/2

 tMatrix['RR','WH', c('WR', 'HR', 'RR')] <- c(1-cM, 1+cM*chM, (cM*(1-
chM)*crM)+(cM*(1-chM)*(1-crM)))/2

 tMatrix['HR','WH', c('WH', 'HH', 'HR', 'WR', 'RR')] <- c((1-cM)*(1-dF), 1+cM*chM + (1-
cM)*dF*dhF, (cM*(1-chM)*crM + 1+cM*chM + (1-cM)*dF*(1-dhF)*drF)+(cM*(1-
chM)*(1-crM) + (1-cM)*dF*(1-dhF)*(1-drF)), (1-cM)*(1-dF),(cM*(1-chM)*crM + (1-
cM)*dF*drF)+(cM*(1-chM)*(1-crM) + (1-cM)*dF*(1-drF)))/4

 #male stuff from female deposition
 tMatrix['WW','HH', 'WH'] <- 1
 tMatrix['WR','HH', c('WH', 'HR')] <- c( 1, 1)/2
 tMatrix['WW','HR', c('WH', 'WR')] <- c( 1, 1)/2
 tMatrix['WR','HR', c('WH', 'WR', 'HR', 'RR')] <- c( 1, 1, 1, 1)/4
```

**Table S9. Fitness, GD, maternal deposition, and GDBI formation parameters used in the MGDrivE modeling for the AeaNosC109$^{GD}$ and AeaZpgC109$^{GD}$ lines.** * = fitness cost defined for GD hemizygotes.

| MGDrivE parameter | description | AeaNosC109$^{GD}$ | AeaZpgC109$^{GD}$ | Reference |
|---|---|---|---|---|
| **betaK** | Daily # of eggs laid per female mosquito | 20.9 | 23.5 | Sanchez *et al.*, 2020 (2); Otero *et al.*, 2006 (3) |
| **tEgg** | Number of days spent in the egg stage | 5 | 5 | Sanchez *et al.*, 2020 (2); Christophers, 1960 (4) |
| **tLarva** | Number of days spent in the larval stage | 9.5 | 9.5 | This paper |
| **tPupa** | Number of days spent in the pupal stage | 2 | 2 | This paper |
| **popGrowth** | Population growth per generation | 1.175 | 1.175 | Sanchez *et al.*, 2020 (1); Simoy, Simoy & Canziani, 2015 (5) |
| **muAD** | Daily death rate for adult mosquitoes | 0.09 | 0.09 | Sanchez *et al.*, 2020 (1); Fay, 1964 (6); Focks *et al.*, 1993 (7); Horsfall, 1955 (8). |
| **cM** | Male homing rate | 0.76 | 0.58 | This paper |
| **chM** | Male correct homing rate | 0.93 | 1 | This paper |
| **crM** | Male gene drive resistance generating rate | 0.07 | 0 | This paper |
| **cF** | Female homing rate | 0.81 | 0.69 | This paper |
| **chF** | Female correct homing rate | 0.93 | 0.97 | This paper |
| **crF** | Female gene drive resistance generating rate | 0.07 | 0.03 | This paper |
| **dF** | Female deposition homing rate | 0.19 | 0.14 | This paper |
| **dhF** | Female deposition correct homing rate | 0 | 0 | This paper |
| **drF** | Female gene drive resistance generating rate as | 1 | 1 | This paper |

| | | | | |
|---|---|---|---|---|
| | result of female deposition | | | |
| **xiM** | Genotype-specific male pupatory success | HW* = 0.13 | NULL | This paper |
| **xiF** | Genotype-specific female pupatory success | HW* = 0.13 | NULL | This paper |
| **s** | Genotype-specific fractional reduction in fertility | 0.1 | 0.1 | This paper |
